# When causes co-occur: Overcoming distortion in multidimensional predictive representation

**DOI:** 10.1101/2025.07.29.667463

**Authors:** Euan Prentis, Akram Bakkour

## Abstract

Predicting how our actions will affect future events is essential for effective behavior. However, learning predictive relationships is not trivial in a multidimensional world where numerous causes bring any one event about. Here we examine (1) how these multidimensional dynamics may distort predictive learning, and (2) how inductive biases may mitigate these harmful effects. We developed a theoretical framework for studying this problem using computational predictive learning models. Model simulations demonstrate how spurious observations arise in such contexts to compound noise in predictive representations and limit the generalizability of learning. We then provide behavioral evidence in human participants for a conceptual inductive bias that constrains these predictive learning dynamics based on the categorical relatedness of causes and outcomes. Together, these results show that conceptual knowledge can shape multidimensional predictive learning, potentially minimizing severe distortions that may arise from complex everyday observations.

## Introduction

To make good decisions, humans must predict how actions will affect future experiences. However, making such predictions is not straightforward in a multidimensional world where numerous causal processes drive how one event leads to the next. The co-occurrence of these causes creates ambiguity about which causes produce which outcomes. Understanding how these causal mappings are disentangled in memory is essential to understanding how humans effectively predict complex futures during decision making.

The ability to make predictive inferences has been long thought to rely on memory representations that encode a structured model of the environment^1–4^. An early formulation of this idea was Edward Tolman’s concept of cognitive maps, that mentally represent the spatial layouts of environments to aid flexible navigation^5^. The neural basis for these representations was later demonstrated through the discovery of place and grid cells in the hippocampal-entorhinal system, which are thought to encode, respectively, specific locations in space and a coordinate system for space^6^. More recent work has extended the apparent role of these representations beyond physical space, showing that similar neural codes organize abstract conceptual information, such as social dimensions^7^, conceptually relevant feature dimensions^8,9^, and a “bird” space that reflects differences in neck and leg length^10^. These codes have also been theorized to implement not just Euclidean maps, but also graph-like representations that capture the relational structure between discrete concepts, states, or events^4,11,12^. Importantly, these relations can be statistical regularities, such as the frequency with which events transition between one another^13,14^. These representations can therefore be used to predict how events unfold into the future.

The reinforcement learning framework formalizes how predictive representations are used in service of goal-directed behavior^15^. In this framework, an agent makes actions to transition between states in an environment, with the aim of occupying rewarding states with desirable features. Predictive representations enable the agent to make inferences about distal states that may be several steps away, facilitating behaviors that will be the most rewarding in the long run. Two influential model classes instantiate this idea. In model-based (MB) learning, a representation of the environment’s transition dynamics is learned and then simulated over at choice to infer distal outcomes. Successor features^16–19^ (SF; a generalization of the successor representation) instead forgoes an explicit model of the transition dynamics, and directly represents each state in terms of the features expected to be encountered over time after entering that state. In both cases, if a state does not contain desirable features but proximal states do, the agent would still choose to approach the state. These models have been found to capture key patterns of predictive learning in behavior^20–22^ and the brain^13,18,22–29^. However, work to date has, to our knowledge, primarily studied how these predictive representations are learned for each state rather than for each of the multiple independent features that compose states. Consequently, it has not been characterized how feature-based representations may be distorted when causal processes, defined over features, co-occur, or whether such distortions are a general property across computational approaches to predictive learning (SF and MB learning).

To illustrate how feature-level causal dynamics may distort representations, consider the following example (Fig. 1A). Suppose two causal processes are defined by feature A1 producing A2 (A1→A2), and feature B1 producing B2 (B1→B2). In a one-dimensional environment, an event might comprise {A1}, which reliably produces event {A2}, allowing A1→A2 to be learned without interference. However, in a multidimensional environment, A1 and B1 may co-occur within a single event {A1, B1}, which reliably produces event {A2, B2}. Since A1→A2 and B1→B2 unfold together, spurious transitions A1⇢B2 and B1⇢A2 will also be observed. When the causal processes consistently co-occur, these spurious and causal observations cannot be disambiguated, leading to faulty or uncertain inference^30,31^. But as we will show, even when causal processes do not consistently co-occur, spurious information can still impair predictive learning. The first aim of the present research is to simulate fully feature-based predictive learning models to demonstrate how these spurious observations may distort inference. We argue that because such spurious observations are inherent to our multidimensional world, predictive representations of real-world experience are inherently prone to distortion.

**Fig. 1:**
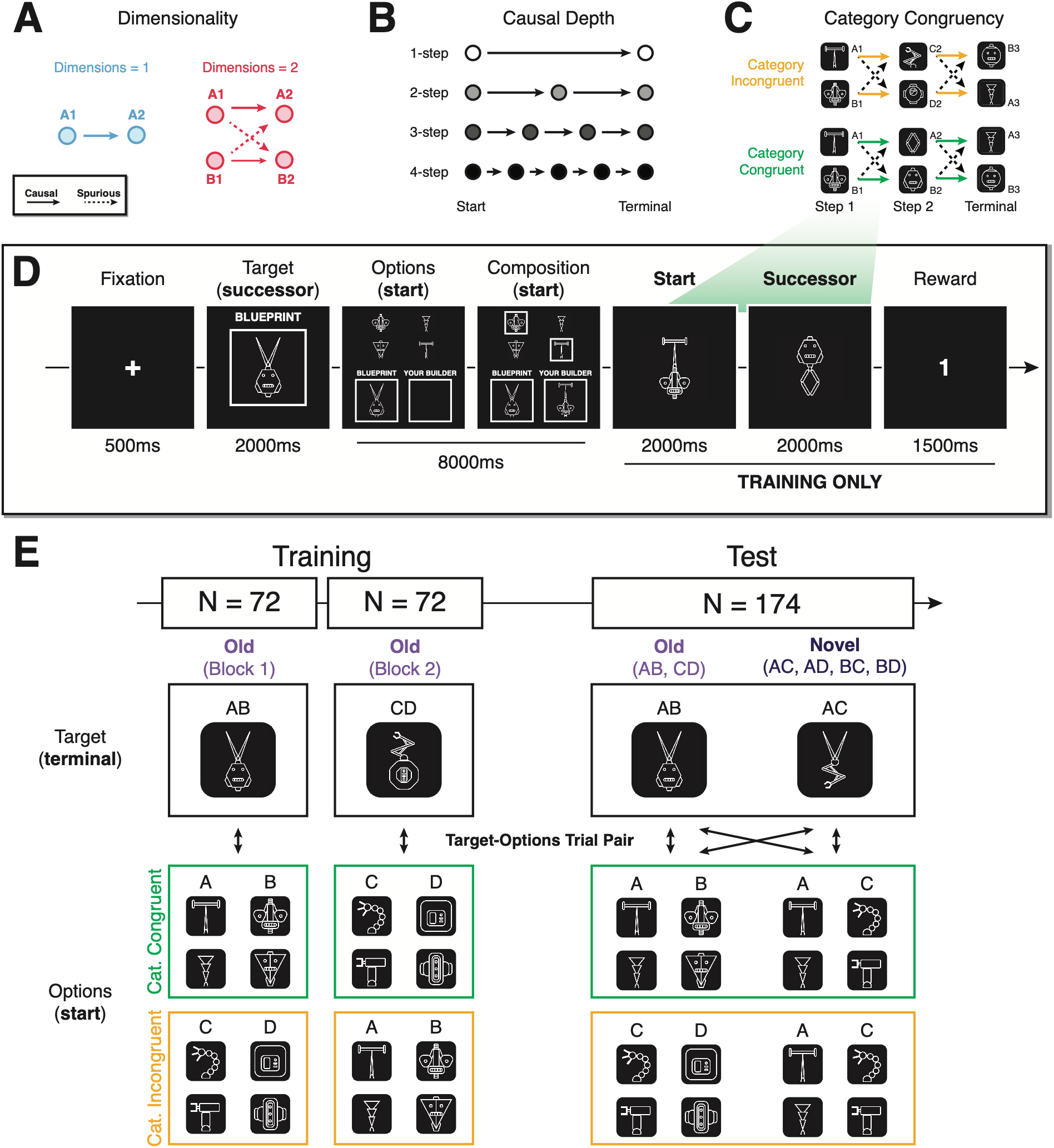
Task design. Simulation environments varied in (**A**) dimensionality and (**B**) causal depth. (**A**) Environments had either one or two features per state. Since A1→A2 and B1→B2 unfold together in the two-dimensional environment, spurious transitions A1⇢B2 and B1⇢A2 are also observed. (**B**) Environments consisted of either (light) 1-, 2-, 3-, or 4-step (dark) causal chains, beginning with *f^START^* and ending with *f^TERMINAL^*. (**C**) Example transition structures in the category-incongruent and congruent conditions. Causal transitions (solid arrows) are defined between step 1 and step 2, and step 2 and terminal features. Although the causal chains are multi-step, only one step is observed per trial of the human experiment. The co-occurrence of features generates spurious observations (dashed arrows). However, whereas all transitions are defined out-of-category in the category-incongruent condition, the causal transitions are defined within category in the category-congruent condition (e.g., head→head). Thus, a conceptual bias would be expressed as upweighted learning of the causal transitions in the congruent relative to incongruent condition. (**D**) Trial procedure (1-step, 2-dimensional environment). Participants are presented with a blueprint for the robot they are to ultimately build. They then compose a builder robot they think will produce the target robot. During training, they then first see the builder (start) robot they composed followed by the final built (successor) robot. Finally, they receive reward commensurate to the number of features that overlap between the successor and target. The example here is from a step-1→step-2 transition in the category-congruent condition, where start and successor features are drawn from the same categories. The top-bottom placement of a robot’s features was randomized within and across trials. For example, here, the start robot was displayed with the antenna on top, while the successor robot was displayed with the antenna on the bottom. (**E**) Feature co-occurrences at training and test, by target and options definition and category-congruency condition in the human experiment (144 unique test trials were shown in the simulations). During training, a subset of feature combinations (AB or CD – old) were presented in each training block. However, at test, both these old combinations and new feature combinations comprising features from across the two training blocks were observed. Moreover, at test, more extensive combinations of old and novel targets and options sets were presented (bidirectional arrows), producing 174 unique trials.

Given this susceptibility to distortion, how do humans learn accurate predictive representations? We suggest that a class of *inductive bias* may attenuate spurious observations during learning. Inductive biases shape learning towards certain interpretations of experience over others^32–38^, and tend to orient towards stable properties of the environment^32,38–40^. This is critically exemplified for causal learning, where prior knowledge has been shown to constrain causal induction, guiding learning towards causal structures that are consistent with existing knowledge^41,42^. Here, we explore whether conceptual knowledge can serve a related role for multidimensional predictive learning. Since conceptual knowledge reflects the statistically regular features and categories of our experiences^43,44^, a conceptual bias may shape learning around these stable properties^32,35,36,40^. Causal processes may be one such property; as the generative components of experience, causal processes are stable across contexts^45^. A conceptual bias may therefore incidentally extract stable causal information while suppressing unstable spurious noise. We test a minimal form of such a bias, which weights predictive learning towards transitions that occur within-category. This bias would favor causal information when causal processes happen to unfold within-category. Thus, the second aim of the present research is to test whether human participants express a conceptual bias on multidimensional predictive learning.

To summarize, the aims of the present research are two-fold. Firstly, we simulate feature-based predictive learning models to demonstrate how spurious observations in multidimensional environments make memory inherently prone to distortion. Secondly, we investigate whether human participants express a conceptual bias on predictive learning that theoretically attenuates the distortive effects of spurious observations.

Results illustrate the crucial role inductive biases play in shaping memory for effective behavior in our complex world.

## Results

### Simulations

To characterize how predictive learning may be inherently prone to distortion in multidimensional environments, we implemented two feature-based predictive learning models: a successor features (SF) learner and a model-based (MB) learner (Methods – Computational Models). These models learned a predictive representation for each feature rather than each state in the environment, and could update representations in parallel as features co-occurred.

The models were simulated on a task in which agents observed simple causal chains defined over features and made compositional predictive inferences to earn reward (Fig. 1). The causal chains could span a single or multiple steps (Fig. 1B; Table S1 – for all definitions), but were always observed to start with feature *f^START^* and end with feature *f^TERMINAL^*. Crucially, these causal chains could co-occur in multidimensional environments, generating spurious observations that may interfere with learning (Fig. 1A&C). On each trial (Fig. 1D – one-step example), the agent was tasked with producing a target terminal state *s^TARGET^* (or ***w***). The agent was then presented a set of start feature options, grouped into a number of categories corresponding to the dimensionality of the environment. The agent selected one feature from each category to compose a start state *s^START^* that may lead to *s^TARGET^*. Then the causal process initiated by *s^START^* and ending with *s^TERMINAL^* was observed. Finally, a reward was paid corresponding to the number of features shared between *s^TERMINAL^* and *s^TARGET^*. The more accurately the agent learned a representation (***M***) that reflected the outcomes of these causal processes, the more reward it earned. Therefore, although reward feedback was graded, the agent never learned reward values through incremental reinforcement learning, meaning the task probed memory for the causal structure rather than reward learning.

Agents completed the task in both one- and two-dimensional environments, wherein states respectively comprised one or two features (Fig. 1A). Since causal processes could not co-occur in the one-dimensional environments but did co-occur in the two-dimensional environments, we could compare learning in the absence versus presence of spurious transitions between these environments. To probe how spurious transitions affected multistep inference, we additionally varied the depth of the causal chains. That is, each agent was simulated in an environment with either one-, two-, three-, or four-step causal chains (Fig. 1B).

We first compare agents’ behavioral performance. A principle of both models is that they should converge on optimal policies in a deterministic environment, irrespective of the environment’s depth. This was demonstrated in the one-dimensional environments (Fig. 2A – Dimensions = 1). Whereas early in training there were some differences across depths in reward earnings, these converged at the same level by the end of training. However, in the two-dimensional environments, performance converged at lower levels in deeper environments (Fig. 2A – Dimensions = 2, green/pink lines). To verify that this decrement in performance was driven by learning about spurious transitions, we simulated versions of the models that were biased towards learning only the causal transitions. Specifically, when incorporating information from the successor state during updating, the causal learners fully discounted information that would be indicative of a spurious transition (that is, *b* = 1; see Methods – Computational Models: Feature-Based Model, Model-Based Learning). Reinforcing that spurious learning drove the performance deficit in two-dimensional environments, the causal-only agents exhibited comparable performance to that of agents in the one-dimensional environments (Fig. 2A – Dimensions = 2, grey lines).

**Fig. 2:**
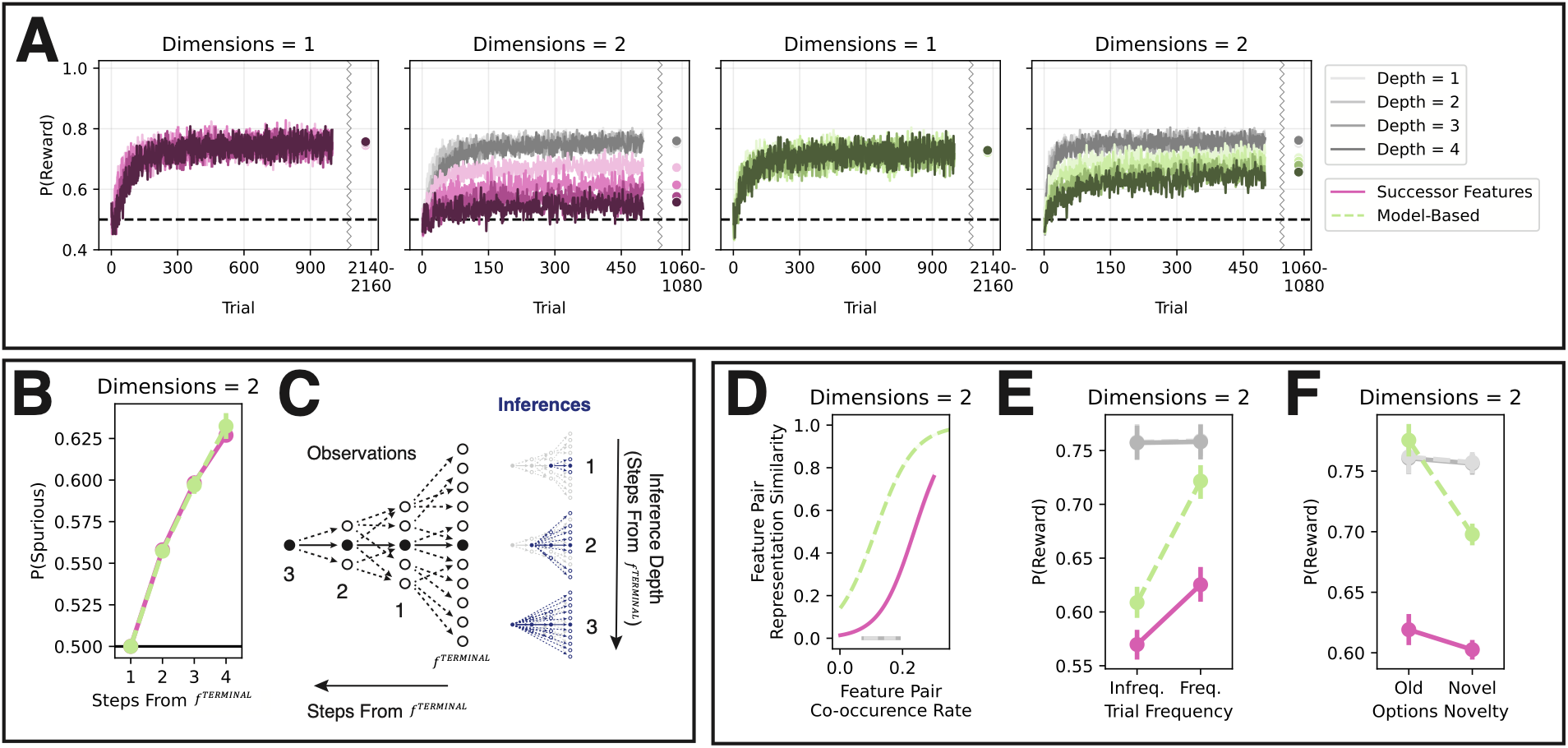
Feature-based model simulations. (**A**) Mean learning curves for the successor features learners (pink lines) and model-based learners (green lines) in one- and two-dimensional environments by causal depth. In the two-dimensional environments, grey lines are causal learners. P(Reward) is the proportion of the maximum reward agents could earn on each trial. (**B**) Proportion of spurious information at inference by model class and the number of steps a feature is from *f^TERMINAL^*. (**C**) Schematic demonstrating how spurious information may compound for deeper features (as shown in **B**). Causal observations (solid lines) are constrained by the deterministic causal processes that generate them, producing simple chains of observations over multiple steps. Conversely, since spurious observations (dashed lines) are not constrained by a mechanism in the environment, they can occur between a more diffuse set of features. This diffusivity expands over multiple steps, producing increasingly less constrained chains of spurious observations. Due to this, features that are deeper in causal chains (more steps away from *f^TERMINAL^*) will compound much more spurious information through these diffuse webs (see **B**). In turn, predictive inferences (red arrows) about deeper features will be noisier and more diffusive, leading to poorer behavioral performance (see **A**, Dimensions = 2). (**D**) The representational similarity of start feature pairs (Pearson’s correlation of 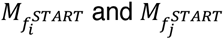, for features 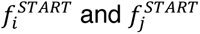) by the rate at which they co-occurred during training in the two-dimensional environments. A separate line is plotted for model class. (**E**) Proportion of reward earned on old test trials by target frequency during training. (**F**) Proportion of reward earned on test trials with just one possible causal and spurious inference by options novelty. All error bars indicate 95% CIs across simulated agents.

Two causal and two spurious transitions were observed on each step in all two-dimensional environments, irrespective of causal depth. Given that this rate of spurious to causal transitions did not vary with causal depth, why did the spurious transitions have a more deleterious impact on performance in the deeper environments? To answer this, we inspected how spurious information accrued in agents’ inferences. Specifically, we isolated how information flowed through memory ***M*** from each feature to all other features by computing predictions for each feature in the task independently (i.e., computing Equations 6&20, defining ***w*** as the identity matrix). We then identified which of these predictions reflected spurious and causal associations, and computed the proportion of spurious to spurious and causal information in these predictions. This showed that spurious information had an outsized effect on inference for deeper features, that were more steps away from *f^TERMINAL^* (Fig. 2B). Thus, even if spurious and causal transitions are observed at the same rate, spurious information can compound through memory for deep features, harming inference (Fig. 2A).

This compounding occurs due to the diffuse nature of spurious associations, that enables spurious noise to propagate widely through memory via SF updating and MB inference (Fig. 2C). Spurious associations are more diffuse than causal associations because they are not constrained by any specific causal mechanism in the environment. That is, an environment with deterministic causal processes will always produce the same causal transition from *f^START^*, while spurious observations will vary simply by what other features *f^START^* co-occurs with across states. Any given feature’s diffuse spurious associations will likely in turn have their own downstream diffuse spurious associations. This broad and deep diffusivity enables spurious information to propagate widely through memory via SF updating and MB inference, particularly compounding for deeper features with more long-run associations. Thus, the processes that enable multistep inference enable spurious noise to compound through memory. Given that real-world experience is driven by deep causal processes, this effect has important implications for predictive learning in naturalistic multidimensional environments.

The distortive effect of spurious transitions on learning reflects not just an injection of noise, but a tuning of representations to specific learning contexts. Spurious transitions in effect act as links between disparate causal processes. If a set of causal processes frequently co-occur, these links are reinforced. Representations of co-occurring features thus converge on common tunings that reflect conjunctive predictions of future features. In short, instead of tuning to the underlying causal processes (e.g., A1→A2, B1→B2), feature representations tune to the learning context (e.g., {A1, B1} → {A2, B2}).

To induce variation in feature co-occurrence frequency, we manipulated the rates at which different target states appeared in the two-dimensional environments (Table S2). Specifically, half the targets occurred twice as frequently as the others, while terminal features occurred at the same rate across these targets. This encouraged consistent compositions on frequent trials, such that certain start features co-occurred more frequently (Fig. S15A). To confirm that the distortions illustrated thus far (Fig. 2A-C) do not depend on this manipulation, we repeated the simulations with all targets occurring at equal rates, such that no combination of features occurred more often than others. These distortions persisted (Fig. S15B&C), indicating they can arise even when features do not consistently co-occur.

To analyze whether representations were tuned based on the frequency with which features co-occurred during learning, for each agent, we characterized how similarly each pair of features were represented by computing the Pearson’s correlation of their corresponding rows in the predictive representation ***M***. Reflecting a tuning to specific feature conjunctions, representational similarity increased as features co-occurred more frequently (pink/green lines in Fig. 2D). The causal learners’ representations did not mirror this effect (grey lines in Fig. 2D), demonstrating that this specificity tuning was driven by the spurious observations.

High specificity tuning had ensuing effects on agents’ inferences. To characterize these inferences after training, agents completed a test phase in which outcomes were hidden to prevent continued learning. Importantly, an extensive set of targets and options were displayed to probe the generalizability of learning (Fig. 1E – akin to category-congruent). Whereas each training half included only “old” targets/options (AB, CD), test also included “novel” targets/options comprising unseen combinations of features (AC, AD, BC, BD). Additionally, all combinations of target and options were shown (e.g., AB-AB, AB-AC, AB-AD, AB-BC, etc.), producing 144 unique trials. Reflecting specificity tuning, more reward was generally earned on: (1) old trials that occurred frequently versus infrequently during training (Fig. 2E); and (2) trials with old versus novel options (Fig. 2F; specifically, this analysis was constrained to trials where only one causal and one spurious inference was possible). The causal learner performed equivalently within both sets of comparisons (grey lines in Fig. 2E&F), again demonstrating that the spurious observations drove both effects.

So far, these simulations demonstrate that spurious observations make predictive learning inherently prone to distortion in multidimensional environments. Specifically, this spurious information makes inference noisier and tunes representations towards idiosyncratic contexts rather than underlying causal processes. Mechanisms that limit the deleterious effects of spurious transitions may thus be essential for effective predictive learning in the real world. But what mechanisms may attenuate this noise?

To start, since spurious information is more diffuse than causal information (Fig. S13), simple mechanisms that *unselectively* erode weak information could attenuate spurious noise while preserving causal predictions. We tested this in the SF Learner, considering three mechanisms: (1) forgetting – ***M*** gradually decays to zero (Equation 16); (2) L1 regularization – the weakest cells in ***M*** are pruned (Equation 17); and (3) non-linear value inference – only the strongest information in ***M*** is used to infer action values. However, simulations of each mechanism showed no benefit for learning, with behavioral performance and the proportion of causal information in ***M*** decreasing with greater forgetting, greater L1 regularization, and more selective non-linear value inference (Fig. S10&11).

Instead, learning may benefit from a more *selective* mechanism that biases memory towards *specific* transitions. Particularly, if this bias accurately shapes learning around causal transitions, spurious noise should be suppressed. In line with this, simulations of an idealized bias defined from the task’s true causal structure (Equations 14&15) show that increasing the degree of bias improved behavioral performance and the proportion of causal information in ***M*** (Fig. S10D&11D). A selective bias towards specific transitions may thus be an essential tool for combating spurious predictive information in multidimensional environments.

But what might direct a selective bias towards causal transitions? We posit that conceptual knowledge could be key to steering this. Conceptual knowledge shapes memory formation^46,47^ and could therefore feasibly also influence which transitions are selected for predictive learning, favoring those that occur within-category. Since categorical structure reflects regularities that are stable across experience^43,48^, conceptual knowledge may incidentally reflect stable causal processes. Resultantly, a conceptual bias may tune learning towards causal associations when category and causal structure align. In the following experiment, we test for a conceptual bias in this way, by minimally manipulating the congruency between categorical and causal structure.

### Human Experiment

To test whether a conceptual bias shapes human predictive learning, 102 participants were recruited to complete a version of the two-step, two-dimensional task used in the simulations (Fig. 1; 144 training trials, 174 test trials). Unlike the simulations, the two-step structure was learned piecemeal, meaning only a step-1→step-2 or step-2→terminal transition was inferred and observed on each trial. Crucially, we also used visual stimuli (robots) with features that had an implied categorical structure (robot parts were drawn from four categories: heads, antennas, arms, bodies). To enforce these categories prior to predictive learning, an encoding phase was completed in which the six robot parts in each category were presented together with their category label (e.g., “arms”). We then varied across two between-subjects conditions whether this categorical structure was *congruent* or *incongruent* with the causal task structure (Fig. 1C; Table S3). If participants used a conceptual bias to direct predictive learning, we hypothesized that this would be expressed as bolstered learning of the causal structure in the category-congruent condition.

### Evidence for a conceptual bias on predictive learning

The experiment’s key marker of a conceptual bias would be upweighted learning of causal relative to spurious transitions in the category-congruent condition. To assess this, we analyzed the extent to which observations of causal versus spurious transitions influenced choice. This involved fitting a Bayesian multinomial logistic regression to each participant’s training data that predicted the composition made on each trial based on recent causal and spurious transitions. On a trial, a participant could have produced one of four possible compositions, which we arbitrarily label 0, 1, 2, 3. Treating composition 0 as the reference class, we modelled the choice of each of the other three compositions *s* relative to composition 0 with a distinct logit model:

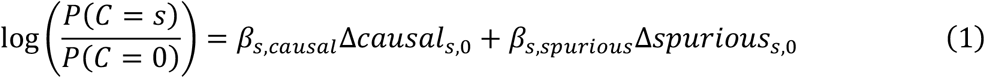

where: (1) Δ*causal_s_*_,0_ is the relative probability that composition *s* versus 0 will produce the target based on recently observed *causal* transitions; and (2) Δ*spurious_s_*_,0_ is the relative probability that composition *s* versus 0 will produce the target based on recently observed *spurious* transitions (see Methods – Causal and Spurious Transition Evidence for more information). Greater *β_s_*_,*causal*_ and *β_s_*_,*spurious*_ fits thus respectively reflect greater uses of causal and spurious learning during inference. Since six coefficients were fit across the three logit models, the three *β_s_*_,*causal*_ and three *β_s_*_,*spurious*_ are metrics that together reflect how much causal versus spurious information drove participants’ behavior in the task.

In line with a conceptual bias, we found that causal relative to spurious transitions had a stronger influence on training choices in the congruent condition (M = 1.847, SD = 0.193, 95% HDI = [1.464, 2.225]; Fig. 3B; Table S6; Fig. S25). Likewise, we observed greater reward earnings in the congruent condition at training (M = 0.143, SD = 0.053, 95% HDI = [0.040, 0.249]; Fig. 3A; Table S7) and test (M = 0.100, SD = 0.019, 95% HDI = [0.063, 0.139]; Fig. S17; Table S8). These results suggest a conceptual bias was leveraged in the congruent condition to better learn the causal predictions and earn more reward.

**Fig. 3:**
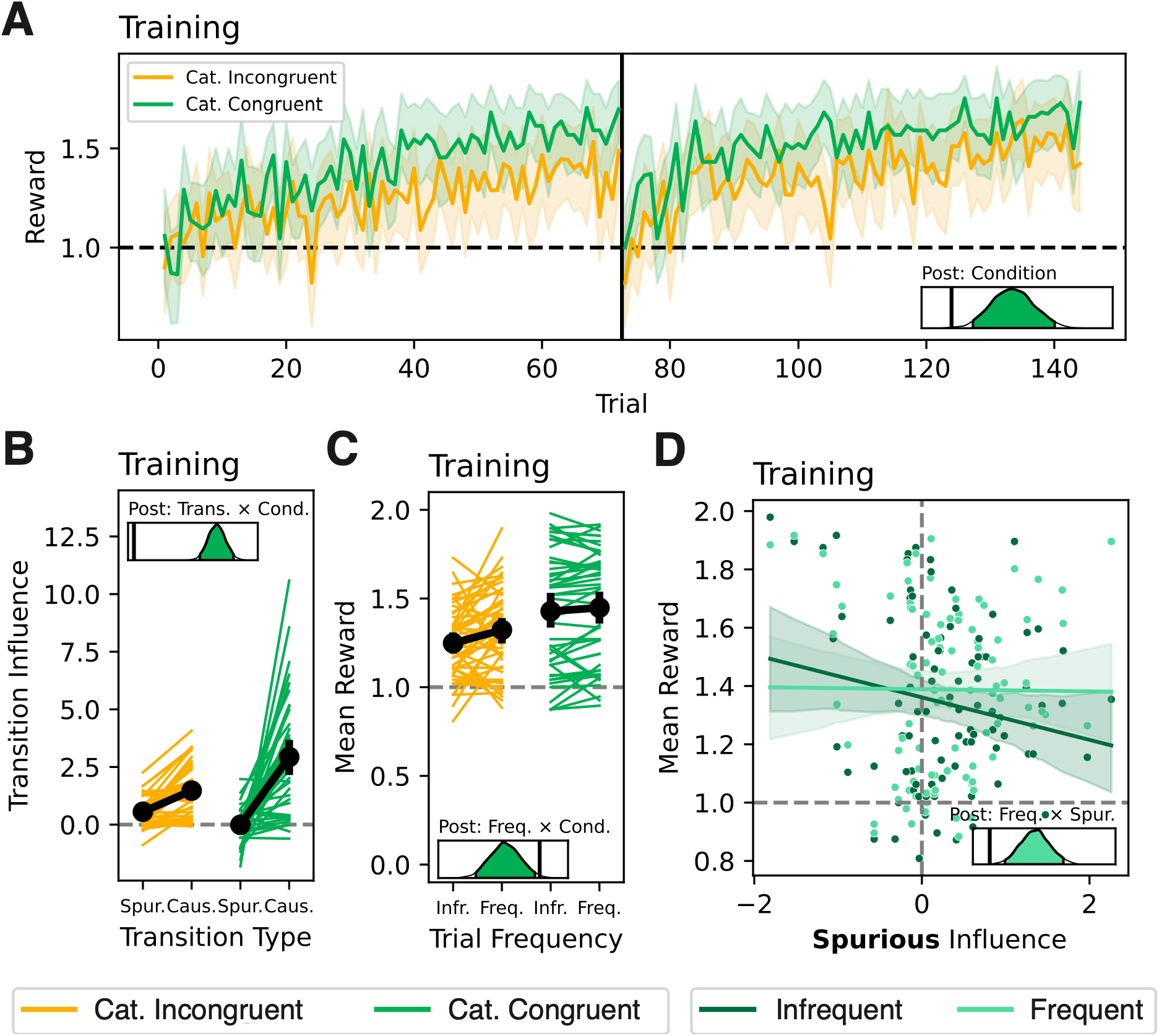
Training Performance. (**A**) Training learning curves by category-congruency condition. (**B**) Spurious and causal transition influence coefficients by category-congruency condition. (**C**) Training reward earnings by training target frequency and category-congruency condition. (**D**) Training reward earnings by spurious transition influence coefficient fit. All error bars are 95% HDIs. Density plots show posteriors from the Bayesian regression models, with black lines indicating 0.

### Conceptual bias supported multi-step inference

The simulations demonstrated that spurious information compounds through memory, producing an outsized harm to deeper multi-step inference (Fig. 2A-C). Since the use of a conceptual bias in the category-congruent condition limited spurious inference (Fig. 3B), it follows that the bias would also improve multi-step inference in the congruent compared to incongruent condition.

To test this, we analyzed whether participants in the congruent condition were better able to make multi-step inferences at test. While during training participants only inferred and observed one-step transitions (options→target: step-1→step-2, step-2→terminal), on 60 test trials with novel options and/or targets, participants inferred two-step transitions (options→target: step-1→terminal). Supporting the hypothesis that the conceptual bias improved multi-step inference, reward earnings on two-step test trials were higher in the congruent than incongruent condition (M = 0.052, SD = 0.018, 95% HDI = [0.018, 0.088], Fig. 4B, Table S9).

**Fig. 4:**
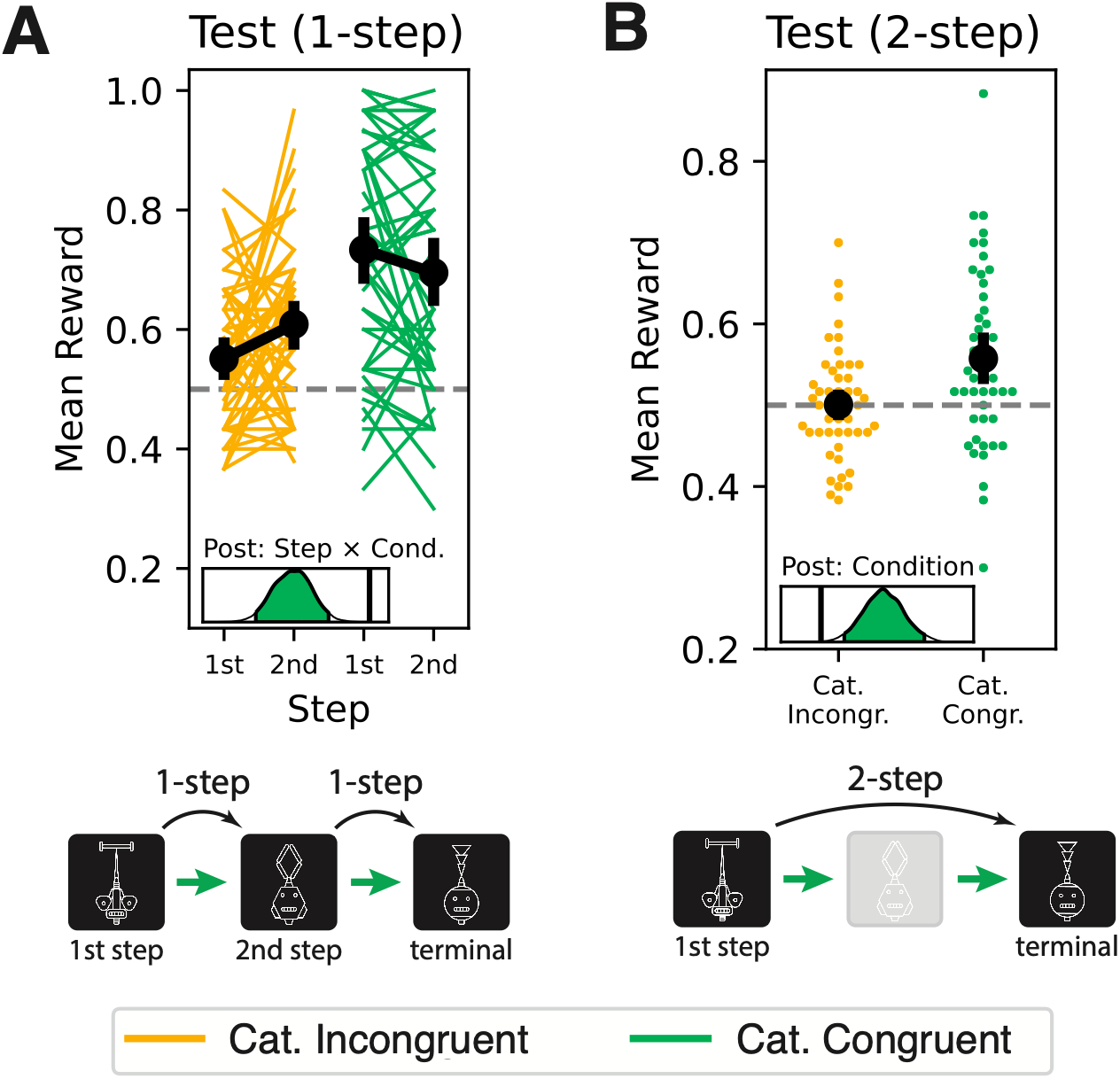
Multi-step inference. (**A**) Mean reward earnings on 1-step test trials by start step and category-congruency condition. (**B**) Mean reward earnings on 2-step test trials by category-congruency condition. All error bars are 95% HDIs. Density plots show posteriors from the Bayesian regression models, with black lines indicating 0.

Multi-step inference may be poorer for deeper features because their representations accrue more spurious noise. If these deeper feature representations are noisier, then even one-step inferences may be poorer for deeper compared to shallower features. We therefore hypothesized that performance would be worse on one-step trials starting from the first versus second step in the incongruent compared to congruent condition. Supporting this hypothesis, reward earnings were lower on first-versus second-step trials in the incongruent relative to the congruent condition (M = -0.094, SD = 0.023, 95% HDI = [-0.139, -0.047], Fig. 4A, Table S10).

Together these results reflect the simulations, suggesting that by attenuating spurious learning the conceptual bias improved multi-step inference.

### Conceptual bias reduced learning specificity

The simulations demonstrated that spurious learning binds features, producing context-specific predictive inferences (Fig. 2D&E). More spurious learning in the category-incongruent condition should therefore produce more specific inferences. In line with this, we found that reward earnings in the congruent compared to incongruent condition differed less by trial frequency (M = -0.063, SD = 0.024, 95% HDI = [-0.110, -0.016]; Fig. 3C; Table S11). To validate these results, we repeated the analysis on a subset of “old” test trials, which had the same targets and feature options from training, only choice outcomes were not displayed. However, we found no interaction between condition and frequency (M = -0.027, SD = 0.056, 95% HDI = [-0.135, 0.085]; Fig. S18, Table S12).

Central to the specificity hypothesis is the rationale that spurious learning tunes representations towards frequent robot transitions. It should therefore be expected that – irrespective of condition – participants whose choices were more strongly influenced by the spurious transitions should also display more sensitivity to the frequency manipulation. An exploratory analysis supported this prediction, showing that participants earned more reward on frequent than infrequent trials when their training choices were overall more strongly influenced by spurious transitions (M = 0.047, SD = 0.013, 95% HDI = [0.023, 0.073]; Fig. 3D, Table S13). Conversely, there was no relationship between causal transition influence and reward earnings on frequent versus infrequent trials (M = 0.007, SD = 0.004, 95% HDI = [-0.0003, 0.014]; Fig. S19). Together, these results demonstrate that higher learning specificity was driven by greater spurious learning.

We can again validate these results by repeating the analysis on old test trials. However, we did not find that test reward earnings differed by training trial frequency and training spurious transition influence (M = 0.023, SD = 0.030, 95% HDI = [-0.035, 0.083]; Fig. S20, Table S14) nor by training trial frequency and training causal transition influence (M = 0.013, SD = 0.008, 95% HDI = [-0.003, 0.030]; Fig. S21).

In all, these results suggest that while attenuated spurious learning in the category-congruent condition reduced learning specificity, this effect did not carry through to test. We speculate that an alternative inference mechanism was used at test that decoupled the relationship between spurious learning and specific inference.

### No evidence the conceptual bias improved generalization

The simulations demonstrated that by tuning learning to frequent state transitions, spurious learning should impair inference not only about infrequent states, but also about entirely novel states (Fig. 2F). A conceptual bias that reduces spurious learning should thus improve generalization to novel states during the test phase.

We evaluated participants’ abilities to make inferences about novel states on trials with novel options and make inferences for novel tasks or goals on trials with novel targets.

We predicted generalization about both would be impaired in the incongruent condition, where spurious learning was unattenuated by the conceptual bias. However, we found no differences in reward earnings by condition and options or target novelty (options: M = 0.014, SD = 0.021, 95% HDI = [-0.028, 0.056], Fig. 5A; target: M = 0.018, SD = 0.021, 95% HDI = [-0.024, 0.059], Fig. 5B; Table S15). Again, we speculate that an alternative inference mechanism may have been used at test to support generalization in the incongruent condition, despite greater spurious learning.

**Fig. 5:**
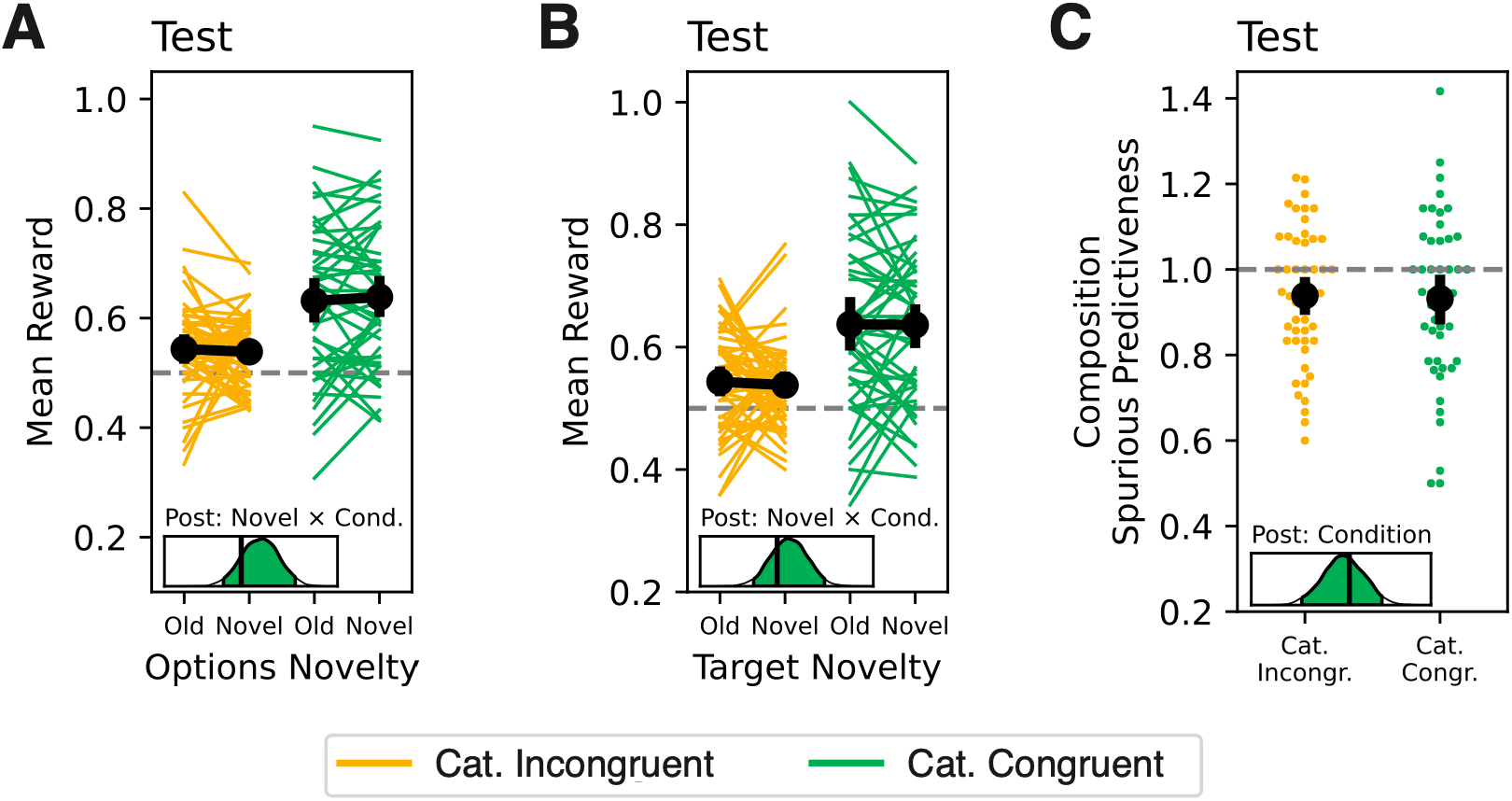
Test Performance. (**A**) Test reward earnings by category-congruency condition and options novelty. (**B**) Test reward earnings by category-congruency condition and target novelty. For **A** and **B**, analyses were restricted to test trials where only one possible causal and spurious inference was possible based on the options and target. (**C**) Composition spurious predictiveness by category-congruency condition on test trials where only spurious inferences were possible based on the options and target. All error bars are 95% HDIs. Density plots show posteriors from the Bayesian regression models, with black lines indicating 0.

We expected generalization to be less accurate in the incongruent condition because greater spurious learning would interfere with causal transition inferences. However, we also predicted that spurious learning could be associated with qualitative differences in the use of spurious information during inference on certain novel trials. Namely, there was a subset of test trials on which only spurious inferences could be made. For example, on a trial in the congruent condition with options AC and target BD, no causal transition and two spurious transitions (A⇢B, C⇢D) would have been observed during training. We hypothesized that greater spurious learning in the category-incongruent condition would encourage participants to actively use spurious information during inference on these trials.

To test this, for each composition, we computed composition spurious predictiveness defined as the number of transitions between the composition and target features that would have been spuriously observed during training. This measure can also be thought of as the reward that would have been received if the causal task structure was defined in terms of these spurious transitions. However, we found no difference in composition spurious predictiveness by condition (M = -0.018, SD = 0.038, 95% HDI = [-0.090, 0.059]; Fig. 5C; Table S16). Intriguingly, when analyzed ignoring condition, composition spurious predictiveness was below-chance (chance = 1, M = 0.944, SD = 0.019, 95% HDI = [0.908, 0.981]). This suggests that participants may have actively avoided making spurious inferences, even if that involved inferring transitions that were never observed during learning.

### Idiosyncratic biases captured by the computational models

Each person’s conceptual knowledge is idiosyncratic, likely producing individual differences in assumed start-successor feature mappings. We should therefore expect idiosyncratic differences in conceptual bias between participants. However, our analyses thus far have assumed that a conceptual bias in the robot task arises solely based on discrete differences between contrived categories (i.e., heads, arms, bodies, antennas) and should be common across participants. We therefore sought to capture individual differences in bias using our computational modelling framework.

The feature-based SF and MB models were fit separately to each participant’s data. Crucially, we fit versions of the models that included a causal-bias parameter (*b* = (0, 1)) that governed the extent to which information indicative of a spurious transition was downweighted during updating (Equations 15&21). Since *b* is agnostic to the underlying memory structure that helps the learner attenuate spurious observations, it can capture individual differences in bias that go beyond the task’s categorical structure.

As an alternative to the selective bias, we additionally fit versions of the SF model that non-selectively eroded weak information in the predictive representation using forgetting, L1 regularization, and non-linear value inference (see Methods, Computational Models, Feature-Based Model). We fit versions of these models that either only included the non-selective mechanism, or additionally included the selective bias.

Lastly, while we have assumed so far that participants learned at the level of features (feature transitions), it was entirely possible for them to learn at the level of feature conjunctions (robot transitions). Since the inductive bias acts on feature-based transition dynamics, it would be invalid to interpret differences in *b* for conjunctive learners. We therefore sought to identify which participants performed feature-based versus conjunctive learning, and verify that the former generally captured behavior better in the task. To do this, we compared the fit of the feature-based models to two *conjunctive learner* models. The basic *conjunctive* model represented each robot as a unique state, acquiring distinct predictive representations for different robots that may share a common feature. The *conjunctive sampler* model learned about robots in an identical way, only it also possessed an inference mechanism that enabled it to retroactively integrate prior learning at choice. The advantage of the latter model is it can generalize learning to novel states, better reflecting participants’ abilities to make inferences about novel options and targets during test.

To analyze individual differences in causal bias, we first needed to identify feature-based learners, for whom *b* could be validly interpreted. We characterized whether participants learned at the level of features or feature conjunctions by comparing the fit of the models (Table S22). For each participant, the best fitting model was identified as that which minimized the Akaike information criterion (*AIC*). A null model was also considered by computing an *AIC* based on the random choice probability for each training and test trial (p = 0.25). Statistically verifying that feature-based learning best explained participants’ choices, a multinomial logistic regression found that a larger proportion of participants (p = 0.63) were better fit by a feature-based model than a conjunctive model (p = 0.29, M = -0.757, SD = 0.220, 95% HDI = [-1.195, -0.347]) or the null model (p = 0.08, M = -2.091, SD = 0.376, 95% HDI = [-2.831, -1.359]). To assess the relative strength of the feature-based and conjunctive fits within each participant, we converted *AIC* values to relative likelihoods, which reflect the plausibility of a given model relative to the alternatives. Since this analysis specifically compared feature-based to conjunctive learning, we identified the best fitting model of each type and computed relative likelihoods between these best fits and the null model. This analysis showed that those best fit by either a feature-based or conjunctive model were strongly best fit by it, as revealed by high relative likelihoods (Fig. 6A, feature: RL = 0.9303; conjunctive: RL = 0.8932; null: RL = 0.6347). This suggests that the conjunctive and feature-based fits reflect real differences in representation, helping us confidently identify feature-based learners.

**Fig. 6:**
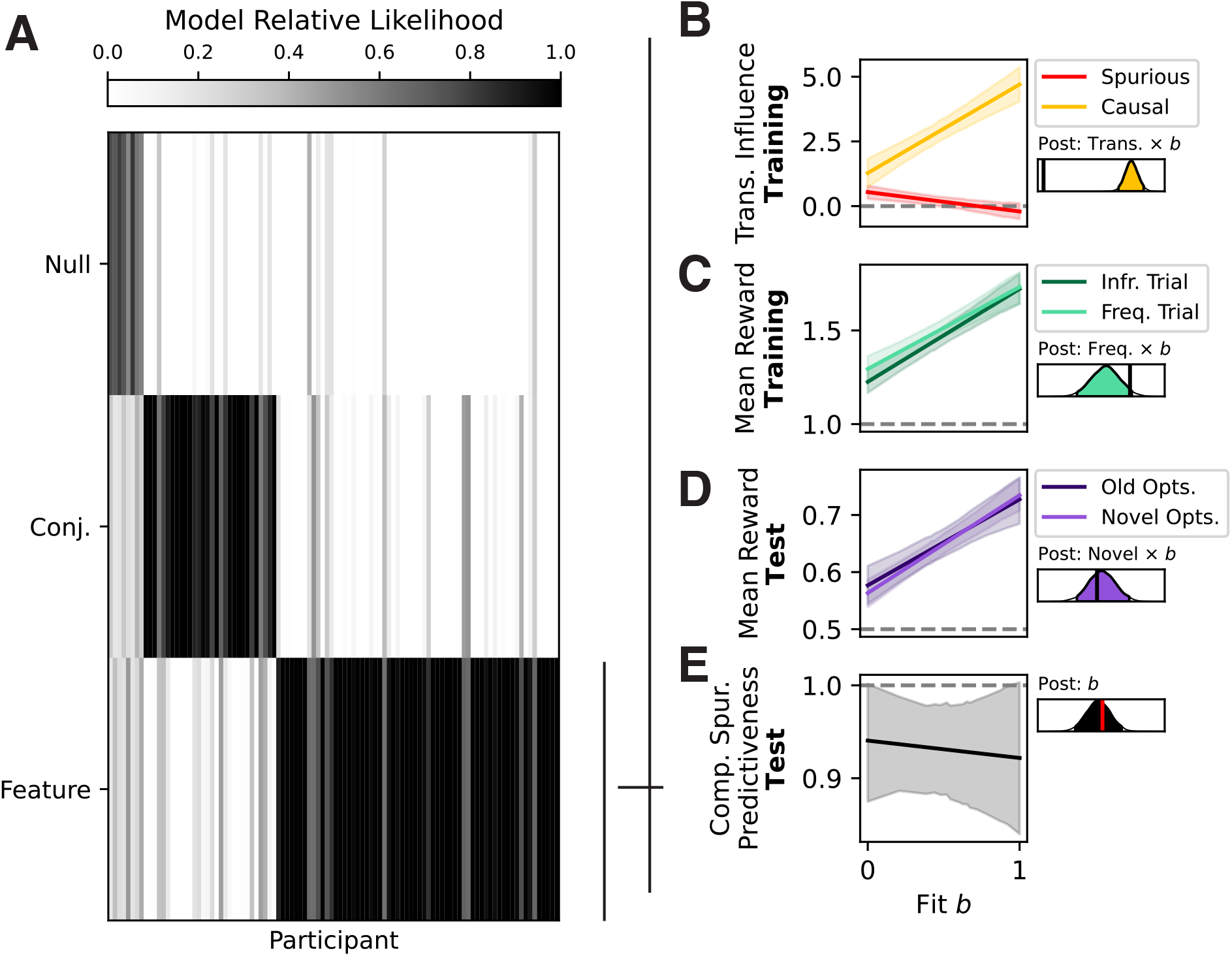
Model-based analyses. (**A**) Relative likelihoods of the best fitting conjunctive and feature-based (and null) computational model by participant. (**B**) Spurious and causal transition influence coefficient fit by *b* fit. (**C**) Training reward earnings on trials with frequent versus infrequent targets by *b* fit. (**D**) Test reward earnings on trials with old versus novel options by *b* fit. (**E**) Composition spurious predictiveness by *b* fit. All error bars are 95% HDIs. Density plots show posteriors from the Bayesian regression models, with black/red lines indicating 0.

Having identified feature-based learners, we could investigate the fit of the causal-bias parameter *b* amongst these participants. If a participant was best fit by a feature-based model that did not include the bias, we assumed *b* = 0. The transition influence analysis demonstrated that participants were biased towards causal information in the category-congruent condition (Fig. 3B). Reflecting this, a beta regression with a logit link function found that *b* was greater in the category-congruent condition (M = 1.166, SD = 0.361, 95% HDI = [0.505, 1.920]; Fig. S22).

Next, we ran a set of exploratory analyses to test whether the fit of *b* captured patterns of behavior in-line with a causal bias.

Firstly, we analyzed whether *b* captured real differences in the attenuation of spurious information. Indeed, a higher *b* fit was associated with greater causal relative to spurious transition influence (M = 4.178, SD = 0.308, 95% HDI = [3.555, 4.755]; Fig. 6B; Table S17).

Next, we analyzed whether a higher *b* fit was associated with lower learning specificity. We observed that reward earnings differed marginally less by trial frequency with a greater *b* fit, during training (M = -0.058, SD = 0.033, 95% HDI = [-0.126, 0.004]; Fig. 6C; Table S18), but did not differ by trial frequency with a greater *b* fit on old test trials (M = - 0.069, SD = 0.078, 95% HDI = [-0.223, 0.081]; Fig. S23; Table S19).

Our analyses of generalization performance by category-congruency condition did not demonstrate differences in choice accuracy on test trials with old versus novel targets or options (Fig. 5A&B). Given that *b* may reflect idiosyncratic biases that differ from the category-congruency manipulation, the fit of *b* may be able to capture differences in generalization at test. However, we did not observe differences in test reward earnings by *b* fit and options or target novelty (options: M = 0.011, SD = 0.031, 95% HDI = [-0.046, 0.073], Fig. 6D; targets: M = -0.020, SD = 0.030, 95% HDI = [-0.079, 0.039], Fig. S24; Table S20).

Finally, we also did not observe differences in the spurious predictiveness of test compositions by category-congruency condition (Fig. 5C). We therefore explored whether the fit of *b* captures individual differences in composition spurious predictiveness. However, we did not observe differences in composition spurious predictiveness by *b* (M = -0.020, SD = 0.057, 95% HDI = [-0.129, 0.091]; Fig. 6E; Table S21).

In all, these model fits largely reflect idiosyncratic differences in causal bias, but, like the model-free results, suggest this may not fully capture how participants generalize at test.

## Discussion

The present work investigated how predictive memory may be distorted in multidimensional environments, and how inductive biases may mitigate these effects. Our simulations illustrate that the spurious observations that arise from multidimensional transition dynamics may both overly tune representations to specific contexts and generate noise that compounds throughout memory. This motivates characterizing inductive biases the brain may leverage to suppress such spurious information. In line with this idea, we found that participants relied on a conceptual bias that selectively shaped learning based on the categorical relatedness of causes and outcomes. We propose that, since conceptual knowledge reflects the stable properties of our experiences^43,48^, this bias tunes learning towards stable causal information in our everyday lives, overcoming harmful distortions in memory.

While participants’ behavior during training largely reflected the predictions of the computational model simulations, some predictions were not supported at test. Specifically, the simulations predicted that a bias towards causal transitions – afforded by a conceptual bias in the category-congruent condition and captured by the computational model – would improve generalization, reduce sensitivity to training conjunction statistics, and limit spurious inference, but we observed none of these effects (Fig. 5, S18, S20, 6D&E). We speculate that the novelty of the test trials prompted participants to adopt a different inference mechanism not captured by our models. People have been shown to adapt how they generalize according to the demands of the environment^49,50^, and selectively generalize dimensions of prior learning that are relevant to a new context^51^. Participants may similarly have employed an alternative inference mechanism at test that limited the influence of spurious associations, producing behavior that deviated from the predictions of the models. In line with this, we observed that spurious inference was below chance on test trials where only spurious and no causal inferences were possible (Fig. 5C), suggesting participants explicitly avoided making spurious inferences. However, this account also has its limits. Performance in the incongruent condition was poor at test irrespective of whether options or targets were old or novel. This suggests that, if employed, an alternative inference mechanism may have been unable to overcome a fundamental degree of noise in the predictive representation that the conceptual bias mitigated during learning in the congruent condition.

The spurious inferences studied here can be considered a case of illusory correlation^52,53^ or illusory causation^54,55^; phenomena whereby a correlation/causation is perceived between two events that does not exist. These phenomena are thought to underlie many maladaptive behaviors, including superstitious thinking^56^, stereotyping^53^, and pseudoscientific medical practices^57^. Prominent theories of illusory correlation/causation suggest that the knowledge, expectations, and attentional biases a person brings to a situation affect how they extract information from the situation, ultimately producing illusory associations^52,58^. In contrast, the present work suggests that conceptual biases may help limit such spurious learning. This discrepancy likely reflects the illusory correlation/causation literature’s focus on explaining real and practically consequential failures of causal reasoning rather than the more elusive situations in which such biases could be beneficial. Conversely, the bias we formalize points towards the true underlying causal model, so our work does not capture how inaccurate biases may actively promote illusory learning. Future work should aim for a more parsimonious account by endowing our models with inaccurate biases and tracking how this affects the emergence of illusory causal models.

Following this line of reasoning, some may debate whether conceptual biases could even be accurate and extract causal information outside of our contrived task setting. This raises the question: can the bias still be beneficial even if it does not direct learning towards causal information? We speculate that it can. Our simulations demonstrated that the high diffusivity of spurious associations enables spurious noise to propagate widely through memory, particularly compounding in deep feature representations (Fig. 2B&C). An inaccurate inductive bias may still help mitigate these effects. By constraining learning to a subset of transitions, the feature associations become less diffusive, and the widespread propagation of spurious noise through memory may be curtailed. Therefore, even if learning is not tuned towards causal information, the memory system as a whole may benefit via the reduced compounding of spurious noise. However, this benefit necessitates that the bias not be too strong. A strong inaccurate bias may limit spurious noise, but also prevent learning of causal transitions not captured by the bias. In the present study, a strong bias was advantageous because the bias was optimal: a maximal bias parameter value and the category-congruent condition both perfectly directed learning towards causal transitions – a design choice intended to encourage the use of the bias, thereby helping us test for it. But in practical real-world contexts, a conceptual bias would rarely, if ever, be optimal. We therefore suggest that this bias is applied softly in practice, guiding learning towards causal information when it aligns with conceptual knowledge, without preventing learning when it does not.

The bias was modelled as a fixed term that applies consistently throughout learning. However, it is possible that participants in the category-congruent condition did not initially apply the bias from the start of training, and only began to do so as they identified the pattern that causal transitions unfold within category. If so, we argue that this would still be an instance of a conceptual bias on predictive learning. The pattern that participants need to detect – that causal transitions are defined within category – depends on participants actively representing the features in terms of the categories. How participants’ conceptual knowledge structures these categories would therefore affect the detection of the pattern. This is in line with the literature on overhypotheses^59^, in which a higher-order regularity (e.g., that causal processes unfold within category) can only be learned over the features a learner already includes in the representation (e.g., including category or conceptual information). Nevertheless, the computational benefits of the bias suggested by the simulations would still apply once the bias emerges, and the structure of a person’s conceptual knowledge may affect how quickly this happens.

Successor features (SF; or the successor representation, originally) was introduced to enable efficient predictive inference in large, multi-step environments without relying on slower model-based (MB) planning^15,17^. Since the computational costs of MB planning scale steeply with the size and depth of the environment, SF has been viewed as especially well-suited for modelling naturalistic inference in real-world environments^3,18^. However, our simulations illustrate that when causal processes co-occur, the advantage of SF over MB in these deeper environments collapses, with spurious information distorting both forms of learning (Fig. 2). This suggests that preserving the advantage of SF in such contexts may depend on useful inductive biases that mitigate these distortions. Notably, the bias implemented here changed not just how SF is learned, but what is learned. Whereas standard (unbiased) learning converges to the analytic successor matrix determined by the environment’s transition structure^17^, biased learning converged to a different representation (Fig. S14), which retained more causal information (Fig. S11D). Thus, learning an agent-specific representation that fundamentally differs to the analytic successor matrix may be beneficial for some learning problems, as exemplified here with co-occurring causes.

The memory distortions characterized by the simulations rest on an assumption of the modelling framework that people, at least sometimes, perform feature-based learning. This is supported by the feature-based model providing the best fit to most participants’ data (63%). Yet still, a sizable percentage (29%) of participants were best fit by a conjunctive model. An extensive literature has long debated whether people learn about multi-feature stimuli as independent elements (feature-based)^60–62^ or as unique configurations (conjunctive)^63,64^. A more recent perspective is that both forms of learning are employed, but their use is mediated by task demands, stimulus properties, and prior experience^65–67^, and depends on different mechanisms in the brain, with configural learning more strongly engaging the hippocampus^68,69^. Our results are consistent with this hybrid perspective, suggesting that the task generally encouraged feature-based learning while a subset of participants learned about stimuli as conjunctions. Future work could investigate why a subset of participants performed conjunctive learning despite the feature-based causal structure. For example, since conjunctive learning does not produce the spurious information that drives these distortions, it may provide a more robust defense against them, albeit at the cost of generalizability.

Prior work has shown that SF captures key characteristics of putative predictive maps in the hippocampus^24,27^. However, the feature-based predictive representations modelled in the present work may be at odds with prominent theories of the hippocampus, which describe it as a conjunctive learning system^11,44,68–71^. Such representations may instead be found in other regions of the medial temporal lobe, for example in the entorhinal cortex, which has been suggested to learn compositional representations that support generalization^1,11,12,51^. On the other hand, the hippocampus itself may have complementary learning systems, with the trisynaptic pathway rapidly encoding conjunctive representations, while the monosynaptic pathway learns the regularities that recur across events^72,73^. Moreover, it has been shown that a biologically plausible neural network that relies on inhibitory competition (thought to support pattern separation in the hippocampus^72,74^) and a bias acquired through Hebbian learning can form generalizable representations that preserve feature information^34^. To identify where feature-based predictive learning may occur in the brain, continuing work is investigating whether the medial temporal lobe encodes the biased representations characterized by our findings. To summarize, the present work introduced a framework for studying the dynamics of predictive learning in multidimensional environments that better reflect the complexity of real-world human experience. We reinforce broad perspectives that effective behavior cannot arise through slow, inefficient associative learning alone, and therefore requires the influence of inductive biases that constrain and accelerate learning^32,35,36,38,40,75^. Particularly, we demonstrated the role existing conceptual knowledge may play in disentangling webs of co-occurring causal information into accurate feature-based predictive representations.

## Methods

### Computational Models

We developed our formal theoretical framework using computational modelling.

An agent’s interaction with its environment was modelled as a Markov Decision Process (MDP), whereby the agent makes actions (*a* ∈ *A*) to step between environment states (*s* ∈ *S*) based on transition function *T*(*s*^′^|*s*, *a*); where the agent transitions from state *s* to successor state *s*^′^ following action *a*. To render this a multidimensional framework, each state comprises *d* features ***φ***(*s*) = [*φ*_1_(*s*), …, *φ_d_*(*s*)], and *T*(*s*^′^|*s*, *a*) emerges from a transition function defined at the feature level:

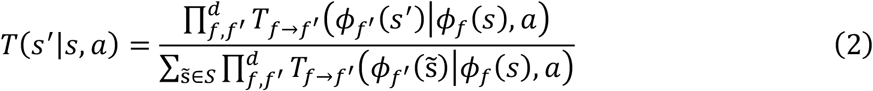

where each feature-level transition function *Tf*→*f*-L*φf*-(*s*^′^)M*φ_f_*(*s*), *a*N specifies the probability of the successor feature *φf*-(*s*^′^), and the denominator normalizes over possible successor states to ensure that ***T*** is a valid transition matrix. This definition of ***T*** assumes that the relationship between observed events is generative, arising from a fundamental set of causal processes that may unfold in parallel. The agent’s goal is instantiated in task vector ***w***, which represents preferences over ***φ***(*s*). The inner product of ***w*** and ***φ***(*s*) defines a reward function ***R***(***w***, *s*), dictating the immediate reward received for entering *s*:

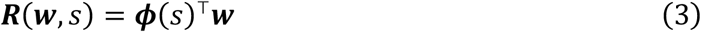

The agent should make actions to navigate towards states with features that will be the most rewarding given the current task ***w***. To infer which actions will give cause to rewarding states, the agent must therefore learn a predictive representation that approximates the transition dynamics defined by ***T***. We model such predictive learning as a successor features (SF) learner^16,17,19^.

The SF model learns a representation of each state *s* in the environment that reflects the quantity of features likely to be encountered upon entering *s* and in subsequent states following *s*. These expectations are represented as row ***M***(*s*) in the successor matrix. We implemented an incremental SF learner that uses a temporal difference rule to learn ***M***(*s*). After state transition *s* → *s*^′^ is observed, the feature vector for the current state ***φ***(*s*) is added to ***M***(*s*). Since this update incorporates the discounted feature predictions of the successor state *s*^′^, the agent gradually learns to predict distant feature visitations. Formally:

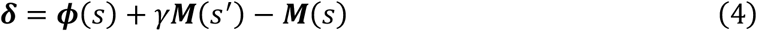

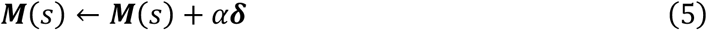

where free parameter *α* = (0, 1) controls the learning rate and free parameter *γ* = [0, 1] controls the discount rate. Since the present task uses one-step terminating transitions, we set *γ* = 1.

To estimate the value of entering *s* given some task ***w***, ***M***(*s*) is used to define a value function ***V***(*w*, *s*) akin to ***R***(*w*, *s*):

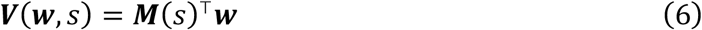

On each trial of the robot task, an agent should compose the robot *s* which has a representation ***M***(*s*) that maximizes ***V***(***w***, *s*) for the trial’s target ***w***. Lastly, a probabilistic choice policy is computed for composing state *s*^∗^ from the set of candidate compositions *C_t_* on trial *t* using a softmax with inverse temperature *β* = (0, ∞):

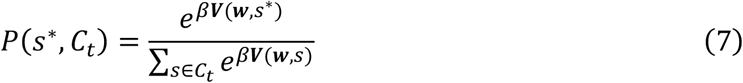

We implemented three forms of this basic SF learner: (1) conjunctive (representing environment states as discrete conjunctions of features); (2) conjunctive sampler (conjunctive representation with a retroactive integration mechanism to generalize to novel states); and (3) feature-based (representing environment states as independent features).

### Conjunctive Model

In contrast to the basic SF learner model described thus far, the conjunctive model represents environment features ***φ***(*s*) in terms of the conjunctive successor state itself. That is, ***φ***(*s*) is a one-hot vector with a length equal to the number of task states (unique robots), and a 1 in the position corresponding to state *s*. Successor matrix ***M*** is therefore square, where the number of rows and columns correspond to the number of unique states. This implementation of SF is akin to the traditional successor representation model.3,17,18

On each trial, the agent constructs an action set that comprises all possible candidate compositions *C_t_* that could be built from the trial’s start feature options. The agent computes ***V***(***w***, *s*^∗^) based on each candidate composition *s*^∗^ in *C_t_*.

### Conjunctive Sampler Model

The conjunctive model suffers from two limitations as a model of human behavior. Firstly, it must compute all possible combinations of features to construct a candidate composition set during inference. This approach may be computationally infeasible in high-dimensional real-world environments where the number of possible feature combinations would likely be intractably large to compute. Secondly, the conjunctive model is unable to generalize learning to make inferences about novel compositions at test. The test phase of the robot task included trials with novel options sets, comprising combinations of feature categories that were not seen during training (AC, AD, BC, BD). Since the conjunctive model would not have learned about the candidate compositions that could be constructed from these options, it could not make inferences about them, and would thus make random choices on novel trials. This does not reflect the behavior of participants, who performed above chance on test trials with novel options (Fig. 5A). Both limitations can be addressed by considering an alternative form of conjunctive model, which represents and learns about states as described, but uses a more flexible inference mechanism to estimate state values at choice.

Previous work has suggested that humans may use a retroactive integration mechanism to sample and combine memories during choice inference^76–84^. This is thought to allow humans to adaptively re-combine prior knowledge into novel configurations to generalize learning to novel problems. We thus implemented a form of retroactive integration that computes weights ***g***(*s*^∗^) over ***M*** based on each represented state’s recency, frequency, and similarity to *s*^∗^ — properties that all drive memory retrieval^85,86^. The estimated value ***V***^∗^(*s*^∗^) is then computed as a weighted sum over ***V***:

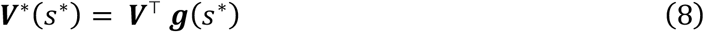

To compute ***g***(*s*^∗^), the agent represents the recency and frequency of states in memory as integer vectors (***g****^REC^*, ***g***^F*RE*G^) with lengths corresponding to the size of *S*. After composition *s* is made on a training trial, a count is added to ***g****^REC^* in all positions except that corresponding to *s*, which is set to zero. A count is also added to the position of *s* in ***g***^F*RE*G^. To allow the similarity computation, the agent stores each observed ***φ***(*s*) in memory. When evaluating *s*^∗^, the similarity *g^SIM^*(*s*^∗^, *s*) of *s*^∗^ to each *s* in memory is calculated as the number of shared features between ***φ***(*s*^∗^) and ***φ***(*s*):

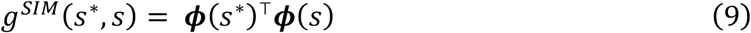

At inference, ***g****^REC^*, ***g***^F*RE*G^, and ***g****^SIM^*(*s*^∗^) are normalized by dividing by their respective maximum values. We then compute ***g***(*s*^∗^) as a weighted sum over these:

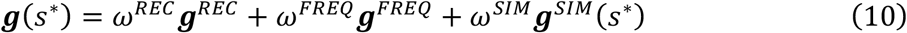

where *ω^REC^*, *ω*^F*RE*G^, and *ω^SIM^* control the degree to which each vector directs memory sampling. To limit the number of free parameters, we treated *ω^SIM^* (*ω^SIM^* = [0, 1]) as the sole free parameter that in turn determined the values of *ω^REC^* and *ω*^F*RE*G^:

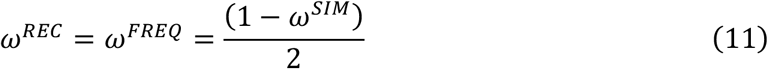

Finally, to capture the extent to which few versus many memories are sampled and integrated over, sampling specificity parameter *ψ* (*ψ* = (1, ∞)) was applied to a normalized ***g***(*s*^∗^):

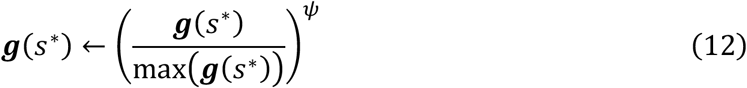

A larger value of *ψ* produced more sampling specificity, directing sampling to the most probable states in memory.

The described retroactive integration mechanism addresses the conjunctive model’s limited ability to generalize. However, it does not address the model’s computationally expensive action set generation mechanism. To account for this, we simply have the model treat the feature options themselves as the action set on each trial, requiring no additional computation. The basic conjunctive model cannot make inferences about these features, since it represents them distinctly to the feature conjunctions it learns about, and does not possess a generalization mechanism to make inferences about them. Since the conjunctive sampler model possesses such a generalization mechanism, it is not limited in this regard. Thus, the conjunctive sampler model can utilize a simpler feature-based action set.

### Feature-Based Model

The feature-based model reflects our primary hypothesis for this study, which is that participants will represent the environment in terms of the independent features that define states. That is, ***M*** is again square, but the rows and columns correspond to independent feature instances rather than conjunctive states. Since a given state transition comprises multiple feature transitions, multiple rows of ***M*** are updated on each trial. This parallelization is implemented by updating the full matrix ***M***, weighted by ***φ***(*s*):

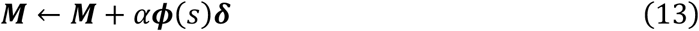

where ***φ***(*s*) is defined as a binarized vector representation of the state’s features, as in the basic SF learner.

The present study investigates an inductive bias towards causal transitions during predictive learning. We implemented this bias via matrix ***B***, where each element ***B****f*,*f*-reflects the probability that features *f* and *f*^′^ are associated. Specifically, we code: ***B****f*,*f*- = 1, if *f* → *f*^′^ is a causal transition; and ***B****f*,*f*- = 1 − *b*, otherwise. Causal-bias parameter *b* (*b* = [0, 1]) dictates the bias magnitude. To apply the relevant rows of the bias given some successor state *s*’, ***B*** is weighted by ***φ***(*s*^′^) during ***M*** updating:

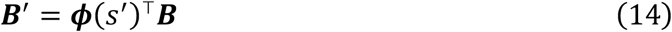

We normalize ***B***^′^ so that rows sum to 1, and then apply the bias on the temporal discounting term

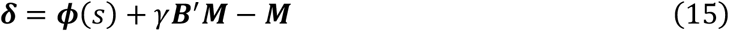

Since causal transitions were defined within categories in the congruent condition, we hypothesized a conceptual bias would direct participants’ learning towards the causal transitions. However, since causal transitions were defined between categories in the incongruent condition, we hypothesized a conceptual bias could not direct learning towards the causal transitions. This pattern of results should be expressed as a higher fit *b* in the congruent condition compared to the incongruent condition.

As an alternative to the selective bias mechanism, we considered three simple mechanisms that unselectively erode weak information in ***M***: (1) forgetting, (2) L1 regularization, and (3) non-linear value inference:

#### Forgetting

Forgetting was implemented via a decay term that degenerates ***M*** towards zero during updating:

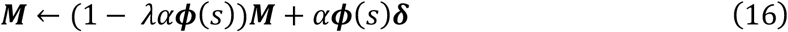

where *λ* = [0, 1] is a decay parameter that dictates the degree of collapse to 0 in rows of ***M*** corresponding to present features ***φ***(*s*). A higher *λ* corresponds to greater forgetting.

#### L1 Regularization

Enforcing a sparse representation may discard weaker spurious noise while preserving stable causal information. L1 regularization enforces sparsity by zeroing cells whose absolute value falls below a threshold. We implement soft thresholding by first shrinking values towards zero according to parameter *λ*_L1_ = [0, ∞), proportional to the learning rate *α*:

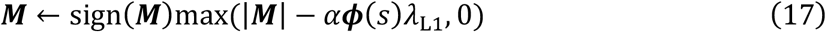

The present state’s features ***φ***(*s*) are included in the threshold to ensure regularization is only applied to the most recently updated rows of ***M***. A greater value of *λ*_L1_ enforced greater sparsity.

#### Non-Linear Value Inference

Sparsity can also be enforced at inference using a non-linear value function. That is, the value ***V***(***w***, *s*) of a state *s* given a task ***w*** is estimated equivalently to Equation 6, ***V***(***w***, *s*) = ***M***u (*s*)^T^***w***, however ***M***u (*s*) is computed by raising ***M***(*s*) to a power *κ* = (0, ∞), normalizing the row ***M***(*s*)***^κ^*** to sum to 1, and then rescaling such that the row’s magnitude matches that of the original representation ***M***(*s*). When *κ* = 1, this recovers Equation 6, and as *κ* increases the sparsity of the predictive inference likewise increases.

### Model-Based Learning

To characterize whether other forms of predictive learning beyond SF are also prone to distortion in multidimensional environments, we implemented an additional predictive learning model: model-based reinforcement learning^15,22^. Whereas the SF learner compresses the environment’s transition dynamics, facilitating rapid multi-step inference at the cost of fine-grained accuracy about single-step transitions, MB instead learns an accurate representation of single-step dynamics while requiring more costly iterative planning to make multi-step inferences.

Specifically, ***M*** is a model of one-step transition dynamics learned using the same update rule as the SF learner (Equation 13), except that the error signal ***δ*** only incorporates information about the successor state’s features:

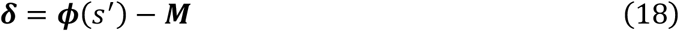

Consequently, ***M*** comes to encode the expected features one-step ahead, rather than the discounted sum of future feature occupancies.

To then infer reward over multiple steps, MB uses a slower planning process whereby feature preferences ***w*** are iteratively propagated through ***M*** by applying the Bellman equation till convergence:

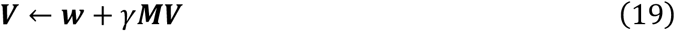

Or, by solving the Bellman equation analytically:

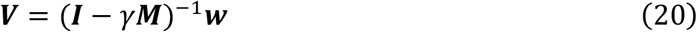

As with the SF learner, we implemented conjunctive, conjunctive sampler, and feature-based versions of MB. By and large, these were equivalent to their respective SF counterparts, differing only by the learning and value-estimation functions described above. The exception to this is that feature-based MB includes an analogous but distinct bias mechanism that applies during ***M*** updating:

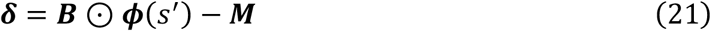

The effect of this is to immediately down-weight one-step transitions that are incongruent with the bias.

### Simulations

In the first set of simulations (Fig. 2), each model was simulated in each task environment 250 times, with random *α* and *β* values. Learning rate *α* was sampled from a uniform distribution bounded *α* = (0, 1). Inverse temperature *β* was sampled as 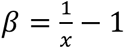, where *x* was drawn from a uniform distribution bounded *x* = (0, 1). For the “regular” learners (pink/green lines in Fig. 2), we set *b* = 0. For the causal learners (grey lines in Fig. 2), we repeated the simulation procedure but set *b* = 1. Agents in the one-dimensional environments were trained for 2160 trials. Agents in the two-dimensional environments were trained for 1080 trials.

### Model Fitting

All computational models (Table S22) were separately fit to each participant’s choice data using maximum likelihood estimation. Given a set of parameter values, the softmax function estimates the probabilities of a participant’s sequence of choices. To fit each model, we computed the negative log likelihood from these probabilities 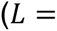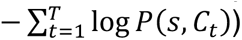, and identified the set of parameter values that minimized this negative log likelihood (using the L-BFGS-B method implemented in python via scipy’s optimize.minimize function). To avoid finding local minima, this fitting procedure was repeated up to 100 times with random parameter initializations. If the negative log likelihood was not minimized further for five successive random initializations, the procedure was terminated.

To account for differences in model complexity during model comparison, we converted *L* to Akaike information criterion (*AIC* = 2(*k* + *L*), where *k* is the number of free parameters), penalizing models with more free parameters. For each participant, the best fitting model was identified as the one that minimized the *AIC*.

## Human Experiment

### Participants

One-hundred and twelve participants were recruited to complete the 90-minute learning task at either the University of Chicago’s campus in Hyde Park, Chicago, or at Mindworks: The Science of Thinking – a behavioral science museum located in downtown Chicago. Ten participants were excluded for exiting the task early either voluntarily or due to a technical error. This left a final sample of 102 participants (mean age = 28.72; standard deviation age = 13.91; 64 female; 26 male; 12 another gender identity). The exclusion criteria and sample size followed a preexisting preregistered experiment (Fig. S1), with the sample size additionally being justified by a power analysis based on those initial results (Supplementary Results, Power Analysis). Participants who completed the study on the University of Chicago campus received either 1.5 points of course credit (N = 31) or $22 plus a fixed bonus of $3 (N = 33). Participants who completed the study at Mindworks (N = 38) received 2200 Mindworks points plus a fixed bonus of 300 points. The Mindworks points were equivalent to $25 in total, and could be exchanged for prizes in the museum. The values of both the monetary and Mindworks bonuses were not revealed until the end of the study. Consent was obtained in accordance with a protocol approved by the University of Chicago’s Institutional Review Board (protocol #: 20-1324).

### Stimuli

The task stimuli were line drawings of robots. Each consisted of two features from four categories (antenna, head, body, arms – randomly mapped to task-relevant labels ABCD). During training, only AB targets appeared in the first block, and only CD targets appeared in the second block. Likewise, only AB/CD options appeared in the first block of the congruent/incongruent conditions, and only CD/AB options appeared in the second block of congruent/incongruent conditions. During test, participants in both conditions were shown robots with all combinations of feature categories (AB, AC, AD, BC, BD, CD). We label robots seen during training as “old” (AB, CD) and new robots only seen at test as “novel” (AC, AD, BC, BD; Fig. 1E). There were six feature instances per category (e.g., six head variations, six antenna variations, etc.). These were assigned to either the 1^st^, 2^nd^, or terminal step of a causal chain (Fig. 1C). Predictive associations were deterministic mappings from 1^st^ to 2^nd^ step instances/items or 2^nd^ to terminal step instances/items (Fig. 1C). In all, there were 24 feature instances (8 1^st^ step, 8 2^nd^ step, 8 terminal) composing 72 items (24 1^st^ step, 24 2^nd^ step, 24 terminal; 24 old; 48 novel).

Every robot had a “flipped” and “upright” version, which only differed by which feature instance was in the top versus bottom position within the stimulus. We randomized which of these two versions was presented for each robot display within and across trials to control for learning and inferences effects based on features’ spatial locations rather than their identities.

### Encoding

To enforce the categories (arms, heads, bodies, antennas) the category-congruency conditions assume, participants were shown all robot parts with their corresponding category label prior to training. That is, for each category, the category label (e.g., “arms”) was shown for 1500ms, followed by a fixation cross for 500ms, and then each of the instances of that category. Each instance was shown for 3000ms, and instances were presented sequentially in a random order. After the four categories had been presented once, the categories were displayed again in a different random order.

### Training

Participants were instructed that they will play a game in which they will act as the owner of a robot factory. Their task is to build robots according to specifications. There are two types of robots: “output” robots (successor items) – the end products the participant is trying to produce; and “builder” robots (start items) – robots that participants themselves compose from sets of builder parts, that in turn produce the output robots. Given the task’s multistep structure, 2^nd^ step robots can be output or builder robots on different trials; 1^st^- vs 2^nd^-step trials, respectively (Fig. 1C, Table S3). On each trial, a “target” output robot is shown. Participants’ goal is to compose the builder that will produce this target. Thus, performance in this task centrally depended on accurate start→successor predictive inferences.

Trials unfolded as follows (Fig. 1D). At the start of each trial, a fixation cross was shown before a target successor item was presented in the center of the screen. Next, an options set of four builder parts (start instances) from two feature categories was shown along with the target. The participant used the cursor to select a feature from each category to compose a start item. As features were selected, the composed item was shown at the bottom of the screen. The two categories were presented on either side of the screen, however which category appeared on which side randomly varied from trial-to-trial. Additionally, the position of each instance within a category randomly varied from trial-to-trial. This spatial randomization ensured that participants’ choices reflected inferences based on features’ identities rather than their spatial locations. After composition, a start→successor sequence was shown, starting with the composed start item, and terminating with the associated successor item produced. Again, we randomized whether an “upright” or “flipped” version of each stimulus was shown to ensure learning was based on the features’ identities rather than their spatial locations. Finally, a point reward was presented. Reward was computed as the number of feature instances that matched between the produced and target terminal items. For these two-feature items, reward values could thus be 0, 1, or 2. To earn a mean reward greater than chance (chance reward = 1), participants needed to learn the start→successor sequences to infer which start robot they should compose to produce the target. To encourage a response on each trial, a -5 reward was given if the participant did not select two features. There were 2 blocks and 72 trials per block, producing 144 training trials in total. The blocks were separated by a 60 second break, and a 60 second break occurred halfway through each block.

We probed the specificity of inference by manipulating the rates at which specific target items were presented (Table S4). Half the targets were presented at twice the rate of the other half of targets. However, features were presented at a uniform rate across these targets. We hypothesized that if participants made consistent inferences about each target, the rates at which they composed start items and observed specific start→successor transitions would converge on the target rates. Our simulations demonstrate that this manipulation should manifest in differential performance for frequent versus infrequent targets only if the spurious transitions are learned (Fig. 2D&E). We thus use differential performance by target frequency as a metric of learning specificity, which we expected to vary with the degree of causal bias.

### Test

Test was the same as training, only participants did not receive feedback for their responses, and they were tested on both one-step (step-1→step-2, step-2→terminal) and two-step (step-1→terminal) inferences for both old and novel targets and options sets. Specifically, there were: 24 old trials that had targets and options both from training (16 one-step: step-1→step-2: 8, step-2→terminal: 8; 8 two-step: step-1→terminal: 8); 30 spurious trials, in which only spurious and not causal inferences could be made; and 120 trials with novel options and/or novel targets. Of these 120 trials, half were one-step (step-1→step-2: 30, step-2→terminal: 30) and half were two-step (step-1→terminal: 60). Because test trials were sampled pseudo-randomly from the set of all possible test trials, the exact split of novel trial types varied slightly across participants: on average, 38.1 trials had novel options only (range = 32–40), 37.9 had novel targets only (range = 32– 40), and 44.0 had both novel options and novel targets (range = 40–52). In total, this produced 174 unique trials. By comparing choice performance on trials with old versus novel options or targets, we could probe the generalizability of participants’ learning. Target frequency was not manipulated at test, and the trial order was randomized. Test was split into two blocks of 87 trials, separated by a 60 second break.

### Category Congruency Condition

A conceptual bias on predictive learning would be expressed as upweighted learning of transitions between categorically-related features, and downweighted learning of transitions between categorically-unrelated features. To test whether a conceptual bias shapes multidimensional predictive learning in humans, we implemented two between-subjects conditions in the robot task (Fig. 1C, n per condition = 51). In the *category-congruent* condition, causal transitions were defined between feature instances from the same category (e.g., start *antennas* always lead to terminal *antennas*). In the *category-incongruent* condition, causal transitions were defined between feature instances from different categories (e.g., start *antennas* may always lead to terminal *arms*). We hypothesized that a conceptual bias would upweight learning of the category-congruent transitions. Thus, there should be better learning of the causal relative to spurious transitions in the congruent compared to the incongruent condition.

### Causal and Spurious Transition Evidence

For the transition influence analysis, causal and spurious transition evidence variables were computed on each trial for each candidate composition as follows. A candidate composition *s* has two features. For each of these features, we found the most recent trial on which it was observed. We then record whether on that past trial the feature transitioned to either (or both) of the target features of the present trial. If a causal transition was observed, a count of 1 was added to the causal transition evidence for *s*. If a spurious transition was observed, a count of 1 was added to the spurious transition evidence for *s*.

For example, *s* may have features A1 and B1, and the target may have features A2 and B4. On the most recent trial A1 was seen, it may have transitioned to a terminal item with features A2 and B4. Thus, the transitions including A1 are: (causal) A1→A2; (spurious) A1⇢B4. Both transitions would also be observed for *s* to the present target, and so a count of 1 is added to the causal and spurious evidence for *s*. On the most recent trial B1 was seen, it may have transitioned to a terminal item with features A2 and B2. Thus, the transitions including B1 are: (causal) B1→B2; (spurious) B1⇢A2. Only the spurious transition would also be observed for *s* to the present target, and so a count of 1 is added only to the spurious evidence for *s*. In all, the counts for *s* would be: causal = 1; spurious = 2. We finally divide these counts by the number of features in the composition (2) to create normalized evidence values: *causal_s_* = 0.5; *spurious_s_* = 1.

Lastly, we convert the transition evidence variables so that they are all relative to reference composition 0:

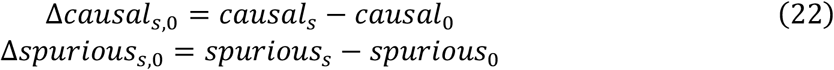

### Bayesian Regression Models

The Bayesian regression models were fit using Markov chain Monte Carlo sampling via the Bambi python package (version 0.14.0)^87^. Four MCMC chains were each run for 4000 to 8000 iterations, with the first 2000 to 4000 iterations of each chain discarded as warm-up. This yielded a total of 8000 to 16000 posterior samples across the four chains. For all models, we assume Bambi’s default weakly informative priors^88^. All model formulas, model fits, and MCMC sampling settings are reported in the supplementary materials.

## Data Availability

Simulation and human participants’ data can be accessed at https://osf.io/nqjy2/?view_only=72ad1546d2df460f9d8b574416c10644

## Stimulus and Task Availability

Task code and stimuli can be accessed at https://github.com/BakkourLabRepo/feat-predict_robot-builder-task-psychopy

## Computational Model Code Availability

Computational modelling code can be accessed at https://github.com/BakkourLabRepo/feat-predict_models

## Analysis Code Availability

Simulation and human experiment analysis code can be accessed at https://github.com/BakkourLabRepo/feat-predict_analysis

## Acknowledgements

We kindly thank Rain Liu for assistance with preliminary analyses during the conceptualization of this work, and Mikul Wyer and Jazmin Velenczei for assistance with data collection. This work was funded by award #2342775 from the U.S. National Science Foundation.

## Contributions

Conceptualization, E.P. and A.B.; Experiment Design, E.P. and A.B.; Computational Modeling, E.P.; Data collection, E.P.; Data Analysis, E.P.; Writing, E.P. and A.B.; Funding, A.B.

