## Supplemental Information for "When causes co-occur: Overcoming distortion in multidimensional predictive representation"

### Supplementary Results

### Online One-Step Experiment

As an initial test for the conceptual bias on multidimensional predictive learning, a preregistered experiment was run in which 100 participants completed the one-step, two-dimensional task used in the simulations (Fig. 2; link to preregistration: https://doi.org/10.17605/OSF.IO/2CY6R). This task was identical to the multi-step task described in the main text, only the causal structure was simplified to one-step and the number of trials was reduced (72 training trials, 144 test trials; Fig. S1).

One-hundred and ten online participants were recruited via CloudResearch's Amazon Mechanical Turk toolkit. Ten were subsequently excluded due to the preregistered criterion of composing a robot on less than 85% of trials. This left a final sample of 100 participants (mean age = 38.92; standard deviation age = 9.95; 31 female; 69 male). Participants received $13 plus a fixed bonus of $3 for completing the 60-minute task. The value of the bonus was not revealed until the end of the study.

The key results of this experiment by and large mirror those of the multi-step experiment in the main text. Participants showed evidence for the conceptual bias on predictive learning, as instrumentalized by a greater influence of causal relative to spurious transitions on training choices in the congruent condition (Fig. S1). Likewise, the conceptual bias limited learning specificity, with reward earnings being more sensitive to conjunctive frequency in the incongruent condition (Fig. S1). A conceptual bias in the congruent condition did not appear to improve generalization to novel options nor targets (Fig. S2). Moreover, computational model fits showed that the feature-based model provided a best quantitative fit to behavior in the task, with the causal-bias parameter $b$ capturing greater causal inference, reduced frequency sensitivity, and reduced composition spurious predictiveness (Fig. S3).

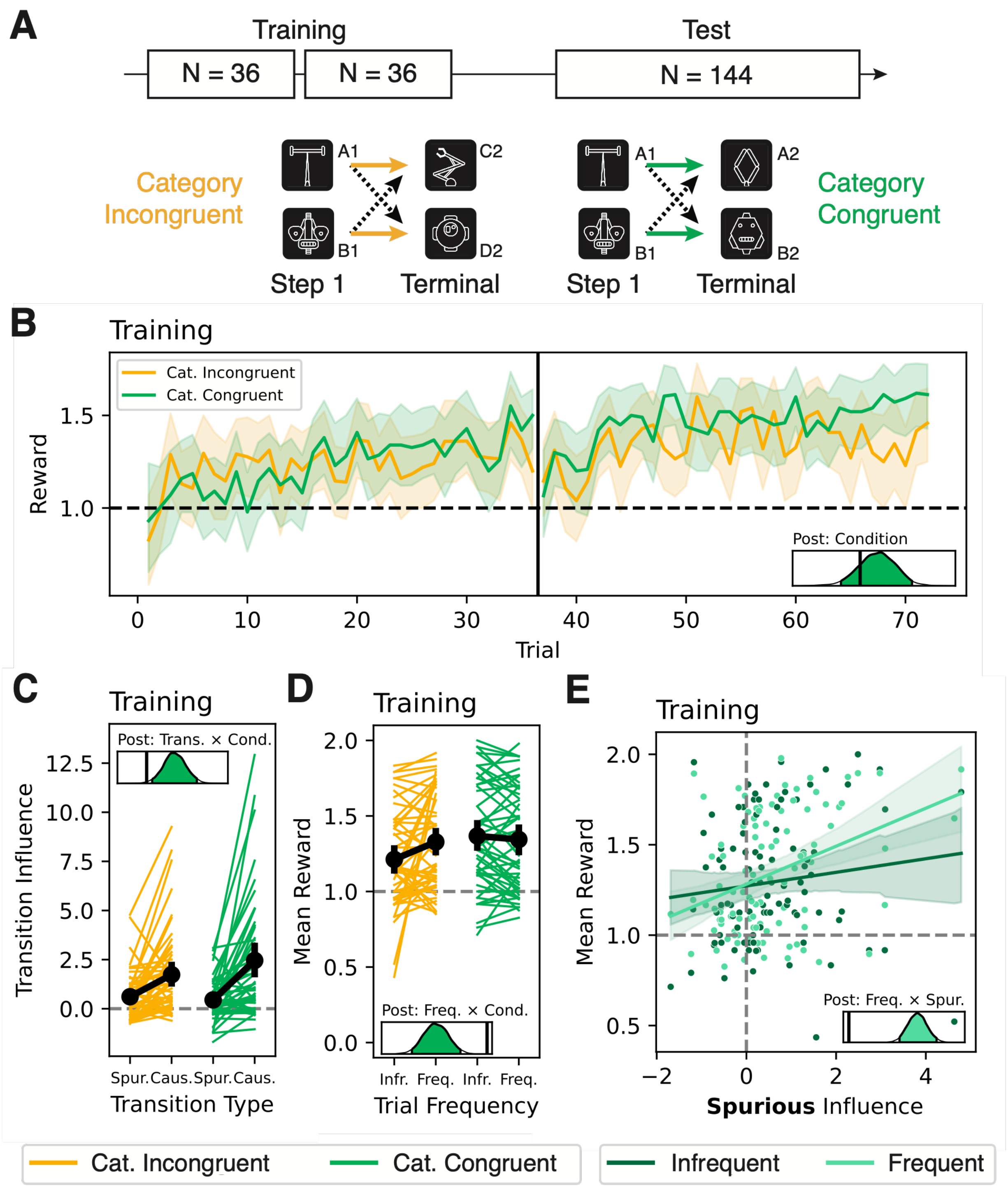


**Figure S1**: **Online one-step experiment – training performance.** (**A**) Task design. The task was identical to the multi-step task (Fig. 1) only the trial numbers were reduced to 72 training trials and 144 test trials, and the task used a single-step causal structure (Table S3). (**B**) Training learning curves by category-congruency condition. (**C**) Spurious and causal transition influence coefficients by category-congruency condition. (**D**) Training reward earnings by training target frequency and category-congruency condition. (**E**) Training reward earnings by spurious transition influence coefficient fit. All error bars are 95% HDIs. Density plots show posteriors from the Bayesian regression models, with black lines indicating 0.

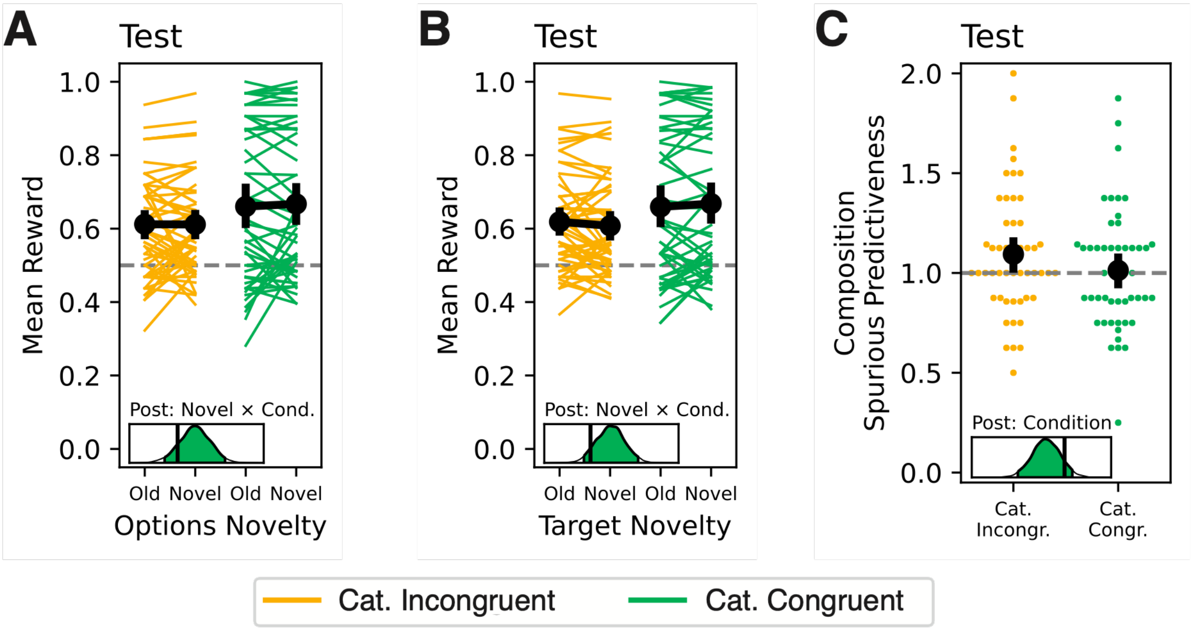


**Figure S2**: **Online one-step experiment – test performance.** (**A**) Test reward earnings by category-congruency condition and options novelty. (**B**) Test reward earnings by category-congruency condition and target novelty. For **A** and **B**, analyses were restricted to test trials where only one possible causal and spurious inference was possible based on the options and target. (**C**) Composition spurious predictiveness by category-congruency condition on test trials where only spurious inferences were possible based on the options and target. All error bars are 95% HDIs. Density plots show posteriors from the Bayesian regression models, with black lines indicating 0.

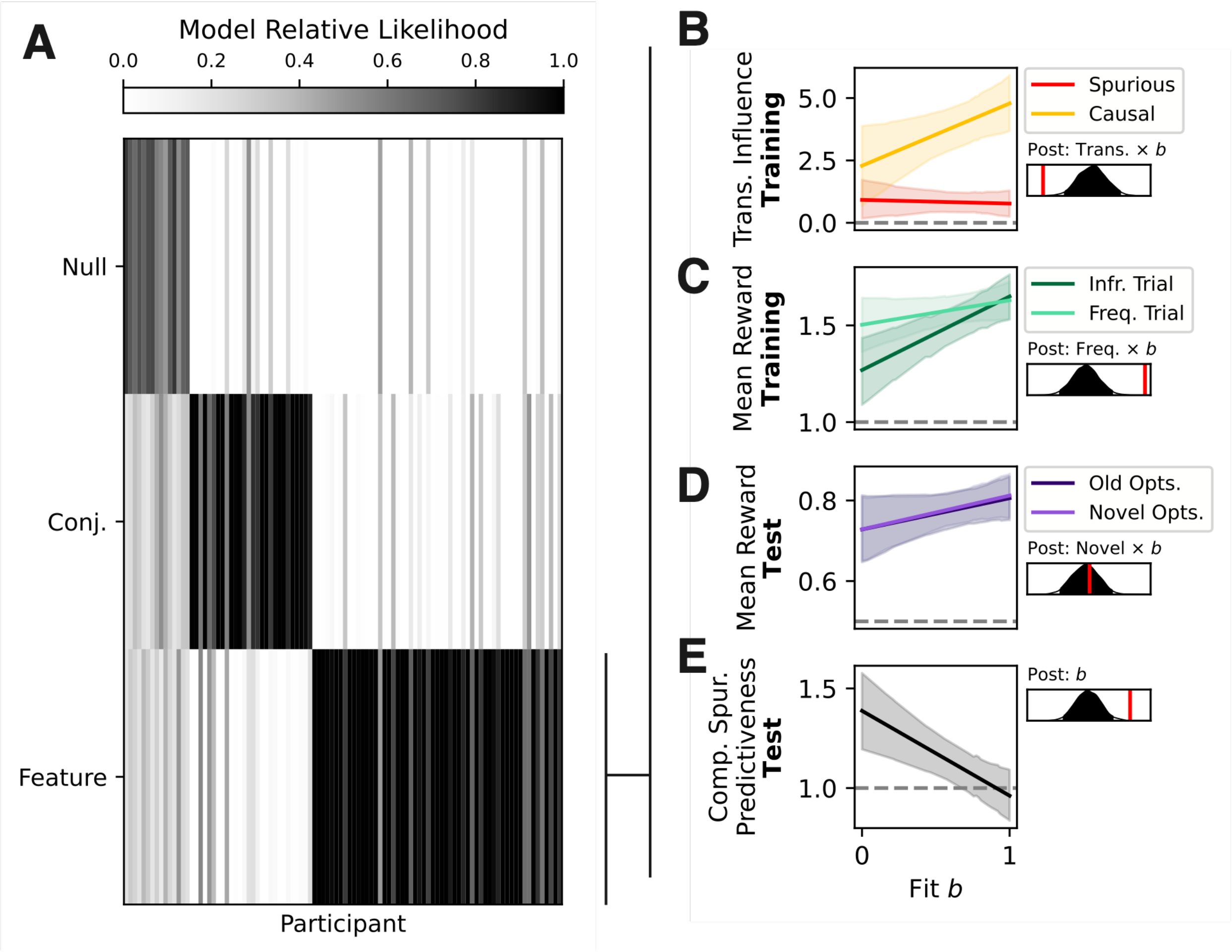


**Figure S3**: **Online one-step experiment – model-based analyses.** (**A**) Relative likelihoods of the best fitting conjunctive and feature-based computational model by participant. (**B**) Spurious and causal transition influence coefficient fit by $b$ fit. (**C**) Training reward earnings on trials with frequent versus infrequent targets by $b$ fit. (**D**) Test reward earnings on trials with old versus novel options by $b$ fit. (**E**) Composition spurious predictiveness by $b$ fit. All error bars are 95% HDIs. Density plots show posteriors from the Bayesian regression models, with red/black lines indicating 0.

### Power Analysis

A target sample size of 100 participants was determined by a power analysis based on the initial one-step experiment (Supplementary materials – online one-step experiment). Our target was to achieve a power greater than 0.8 for the key interaction between transition type (spurious = 0, causal = 1) by category-congruency condition (category-incongruent = 0, category-congruent = 1) in their effect on transition influence coefficient (Fig. S1C; Table S5).

To determine this, we ran 200 simulations of the initial experiment by sampling from the posterior of the second-level transition influence model (Fig. S1C; Table S5). For each simulation, we then fit the second-level model to the simulated data. If the 95% HDI for the interaction between transition type and condition on transition influence exceeded 0 for a given simulation, this was labelled a success. We computed power as the success rate over all simulations.

Using this approach, the initial experiment achieved a power of 0.88 with 100 participants. Therefore, 100 participants is a sufficient sample size to achieve a target power of 0.8, while providing a degree of margin of error should the effect size shrink somewhat during replication. However, to further shield against a deflation in effect size, we additionally doubled the number of training trials the first-level transition analysis is based on from 72 trials in the initial experiment to 144 trials in the new experiment.

### Computational Model Recovery

To verify that the computational models (Table S22) could be distinguished based on fits to behavioral data, we simulated each model in the robot task environment.

Each model was simulated 1000 times with random parameter values (parameters: Table S23). All but two parameters were sampled from a uniform distribution (bounded (0, 0.5) for $\lambda_{\mathrm{L1}}$, (0, 5) for $\kappa$, and (0, 1) for the remaining parameters). The inverse temperature $\beta$ was sampled as $\beta = \frac{1}{x} - 1$, where $x$ was drawn from a uniform distribution bounded (0, 1). Likewise, $\psi$ was sampled as $\psi = \frac{1}{x}$, where $x$ was drawn from a uniform distribution bounded (0, 1). Each model was simulated on trials completed by participants in the multi-step experiment (144 training trials, 174 test trials). All models were then fit to each simulated dataset following the procedure detailed in Methods – Model Fitting. A null model that assumed random choice on each trial was also considered.

To start, we compared the recovery of the feature-based learner, conjunctive learner, and conjunctive sampler learner models across both the successor features and model-based learners. This analysis found that 52%/64% of agents simulated with the feature-based SF/MB model were best fit by the feature-based SF/MB model, 56%/60% of agents simulated with the conjunctive SF/MB model were best fit by the conjunctive SF/MB model, and 34%/37% of agents simulated with the conjunctive sampler SF/MB model were best fit by the conjunctive sampler SF/MB model. Most agents not best fit by the model they were simulated with were best fit by the null model (Fig. S4). Lastly, the SF and MB models were somewhat confusable within the feature-based, conjunctive, and conjunctive sampler classes, reflecting that SF and MB learning produce qualitatively comparable behavior in the task (Fig. 2). Together, these results indicate that the computational models were able to distinguish feature-based and conjunctive representation in the robot task.

Next, we analyzed how differentiable a selective bias was from unselective forgetting, L1 regularization, and non-linear value inference by comparing the recovery of the feature-based SF model between these mechanisms. While the absolute recovery rate between these model instantiations was poor (bias: 47%; forgetting: 21%; L1 regularization: 21%; non-linear value inference: 38%), we note that the bias model was least confusable with the other mechanisms, and most agents not best fit by the model they were simulated with were best fit by the null model (Fig. S5). This indicates that the low absolute recovery rate was driven by simulating in a parameter range that produced noisy behavior. Indeed, when model recovery is assessed across simulated parameter values (Fig. S6), increasing $\beta$ (producing less noisy decision behavior) dramatically improved model recovery. We can therefore conclude that the selective bias' effect on behavior is differentiable from forgetting, L1 regularization, and non-linear value inference.

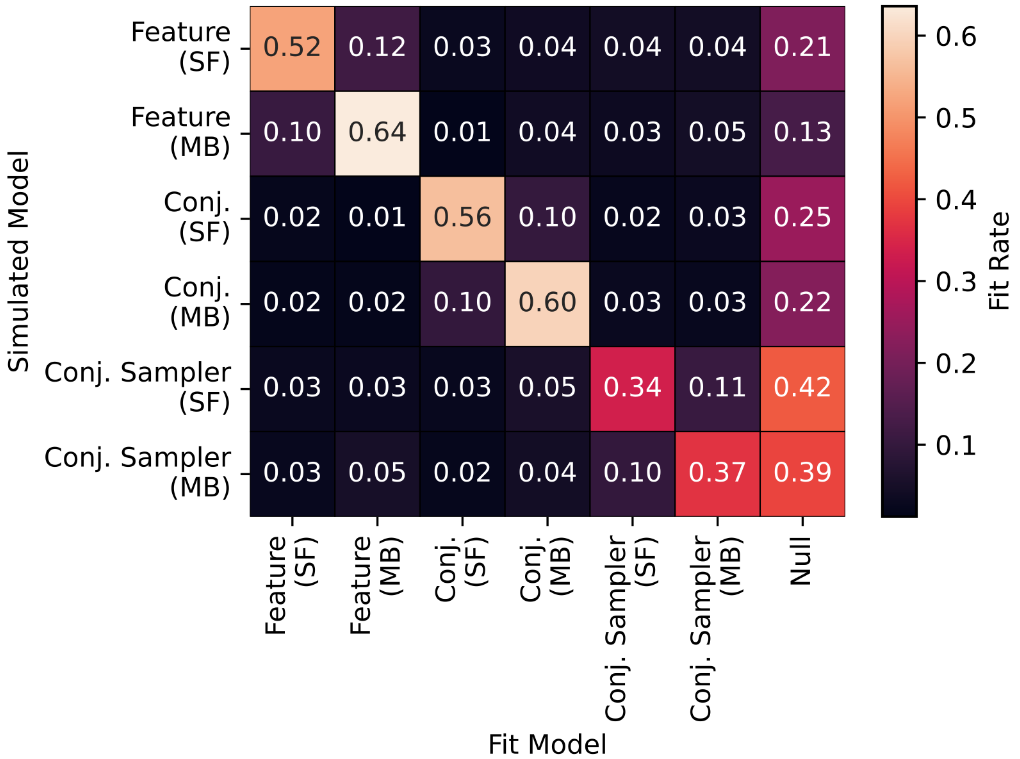


**Figure S4: Model recovery by representation class.** Proportion of best fits by each of the successor features and model-based learner models to each simulated model's data. Simulated models were generally best fit by themselves, with the diagonal being well above the chance rate of 0.143. The successor features and model-based learners were mildly confusable within each representation class (feature-based, conjunctive, and conjunctive sampler) but by and large were separable. However, note that while these model fits were quantitatively separable, the feature-based models produced qualitatively similar simulated behavior (Fig. 2) and the bias parameter $b$ was highly confusable across successor features and model-based learning (Fig. S7).

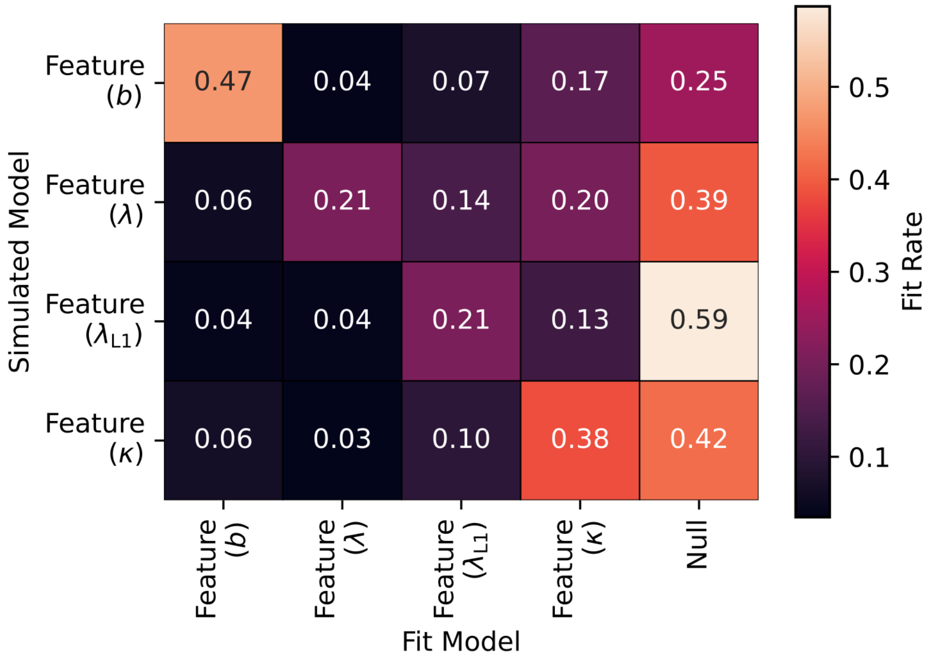


**Figure S5**: **Model recovery for alternative feature-based successor features mechanisms.** Proportion of best fits by each feature-based SF model to each simulated feature-based SF model's data. Simulated models were generally best fit by themselves or the null model. The large proportion of null fits is likely because many combinations of parameters produced noisy simulated behavior. Recovery rates improved as noisy behavior was suppressed (higher simulated $\beta$; Fig. S6).

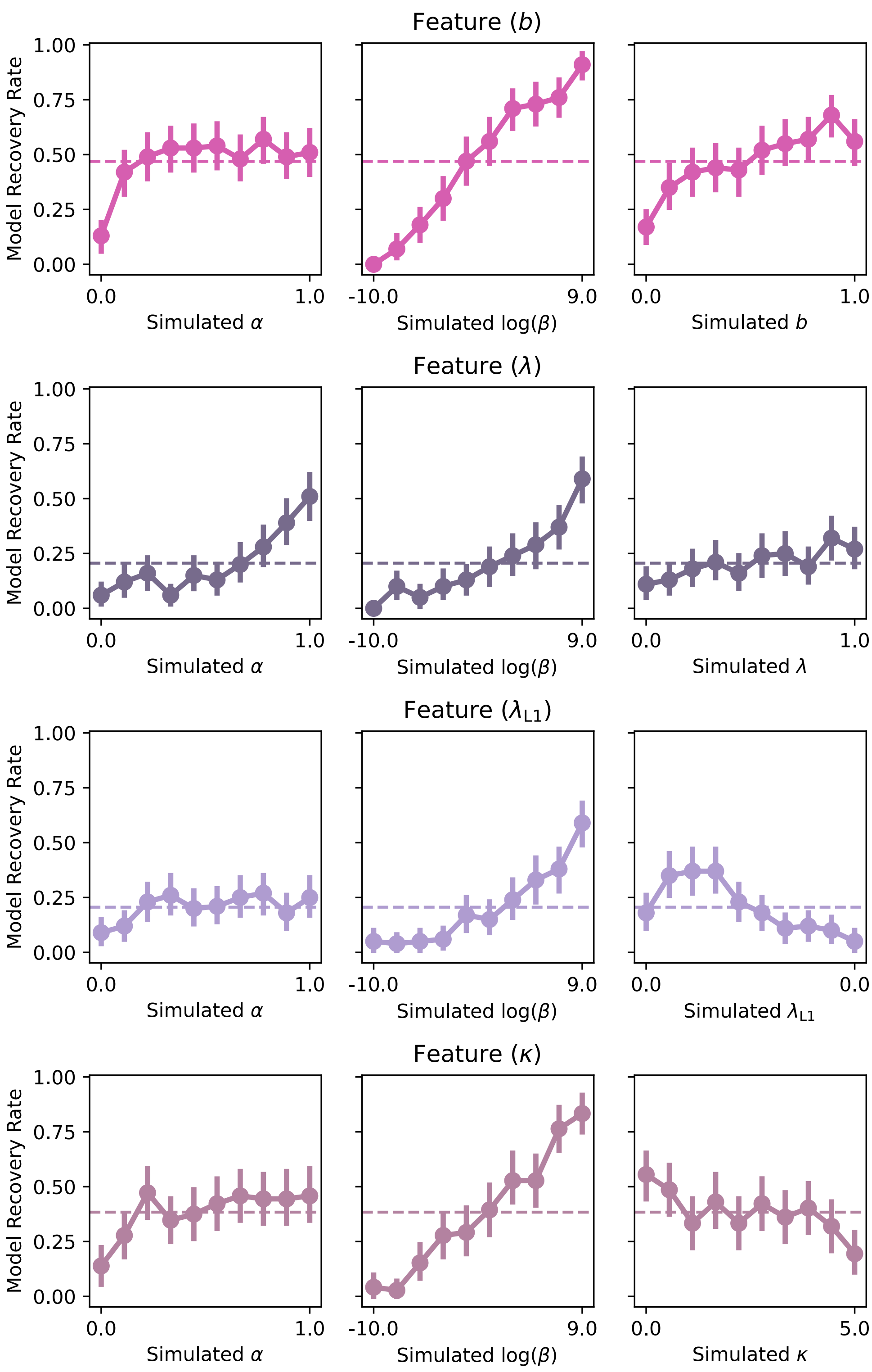


**Figure S6**: **Model recovery within feature-based successor features learning by simulated parameter value.** More deterministic behavior (higher $\beta$) improved model recovery substantially.

### Causal-Bias Parameter Recovery

To verify that the feature-based model's causal-bias parameter $b$ was identifiable based on fits to behavioral data, we evaluated the parameter fits from the model recovery analysis (see Computational Model Recovery). Specifically, we investigated the parameter fits of all feature-based models to agents simulated with the feature-based models.

Firstly, we computed the Pearson's correlation of the simulated and fit $b$. Indicating recoverability within both the SF and MB learners, there was a moderate positive relationship between the simulated and fit $b$ (SF: $r = 0.40$, $p < 0.0001$, Fig. S7A; MB: $r = 0.51$, $p < 0.0001$, Fig. S7C). While the absolute recovery rates of $b$ were not strong, we note that a large proportion (around 13-21%) of simulated SF and MB learners were best fit by the null model (Fig. S4), suggesting that many parameter combinations simply produce noisy simulated behavior rather than $b$ itself being unidentifiable. Indeed, when $b$ recovery is assessed across simulated parameter values (Fig. S8), increasing $\beta$ (producing less noisy decision behavior) dramatically improved $b$ recovery.

Next, we analyzed to what extent $b$ was confusable across the SF and MB learners. Namely, we computed the correlation of $b$ simulated with the SF learner with the $b$ fit with the MB learner, and vice versa. Indicating that these two instantiations of feature-based predictive learning reflect a common bias mechanism, moderate positive correlations showed that $b$ was confusable across the model classes at a similar rate to recovery within each class (simulated-fit – SF-MB: $r = 0.35$, $p < 0.0001$, Fig. S7B; MB-SF: $r = 0.42$, $p < 0.0001$, Fig. S7D). Thus, when interpreting $b$ fits to participants' data, we collapse across the SF and MB model classes and select $b$ from the best fitting model.

To verify that the fit of $b$ was not capturing differences in learning rate $\alpha$ or temperature $\beta$ (i.e., how deterministic choice was), we next computed the Pearson's correlation of the fit $b$ with the fit $\alpha$ and $\beta$. This showed no relationship between the fits of $\alpha$ and $b$ (SF: $r = 0.00$, $p = 0.9535$, Fig. S7E; MB: $r = -0.01$, $p = 0.6985$, Fig. S7G) nor $\beta$ and $b$ for the SF model ($r = 0.01$, $p = 0.7124$, Fig. S7F; $\beta$ were log transformed). There was a small positive correlation between $\beta$ and $b$ for the MB model ($r = 0.10$, $p < 0.001$, Fig. S7H; $\beta$ were log transformed).

Lastly, we compared within SF learner models to what extent $b$ captured and was captured by the forgetting ($\lambda$), L1 regularization ($\lambda_{\mathrm{L1}}$), and non-linear value inference ($\kappa$) parameters. Specifically, we computed the Pearson's correlation of the simulated $b$ within the bias model with the fit $\lambda$, $\lambda_{\mathrm{L1}}$, and $\kappa$ within the forgetting, L1 regularization, and non-linear value inference models, respectively. This revealed negligible negative correlations between simulated $b$ and fit $\lambda$ ($r = -0.1$, $p = 0.0011$; Fig. S9A) and log-transformed fit $\kappa$ ($r = -0.06$, $p = 0.0437$; Fig. S9C), and a small positive correlation between simulated $b$ and log-transformed fit $\lambda_{\mathrm{L1}}$ ($r = 0.19$, $p < 0.0001$; Fig. S9B). The inverse analysis showed similar effects, with no correlation between *fit* $b$ and *simulated* $\lambda$ ($r = 0.02$, $p = 0.5555$; Fig.

S9D) and $\kappa$ ($r$ = 0.07, $p$ = 0.056; Fig. S9F), and a negligible positive relationship between *fit* $b$ and *simulated* $\lambda_{\mathrm{L1}}$ ($r$ = 0.1, $p$ = 0.0010; Fig. S9E). Together, results reinforce the model recovery analysis, indicating that the selective bias' effect on behavior is differentiable from forgetting, L1 regularization, and non-linear value inference.

Together, these results indicate that $b$ was identifiable in the robot task.

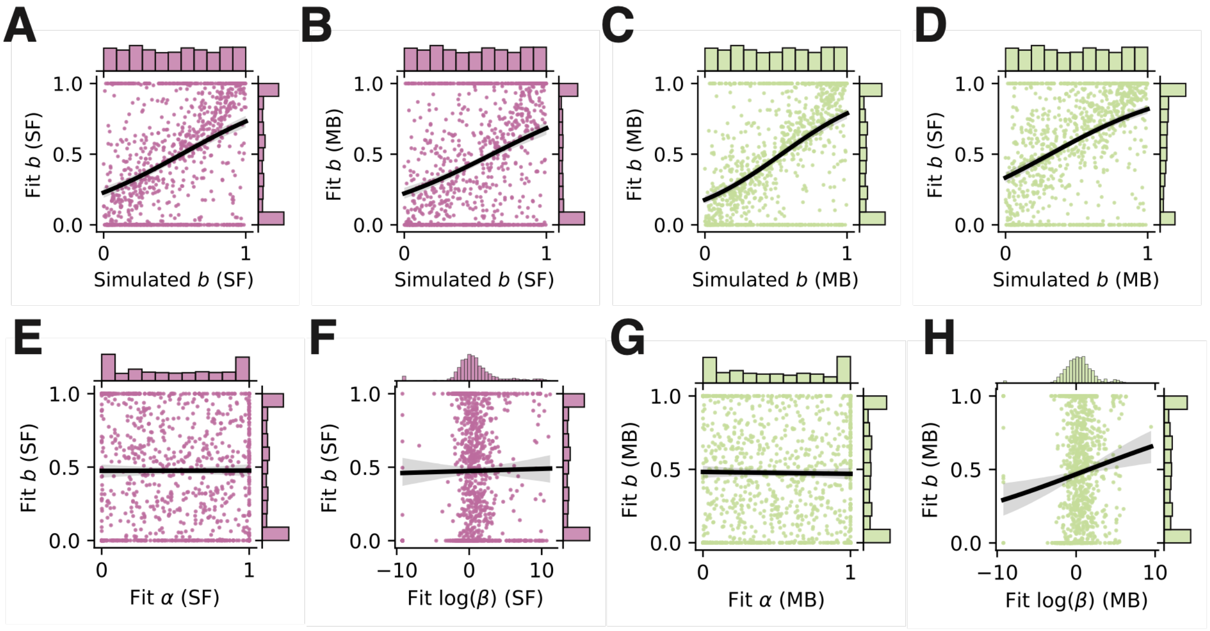


**Figure S7: Feature-based model $b$ recovery analyses.** The $b$ parameter is similarly recoverable whether learning is simulated with successor features and fit with successor features (**A**) or model-based learning (**B**), or simulated with model-based learning and fit with model-based learning (**C**) or successor-features (**D**). Fit $b$ values by fit $\alpha$ within successor-features (**E**) and model-based learning (**G**). Fit $b$ values by log-transformed fit $\beta$ within successor-features (**F**) and model-based learning (**H**).

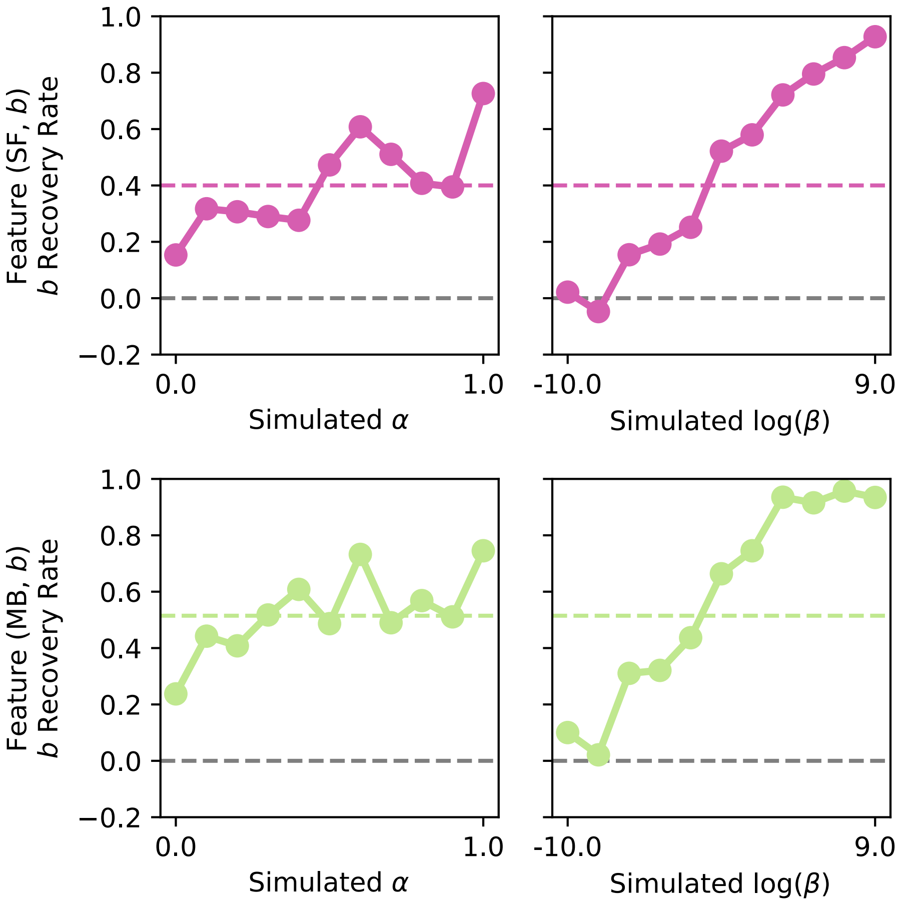


**Figure S8: Feature-based model $b$ recovery analyses by simulated parameter value.** More deterministic behavior (higher $\beta$) improved $b$ recovery substantially for both successor features and model-based learners.

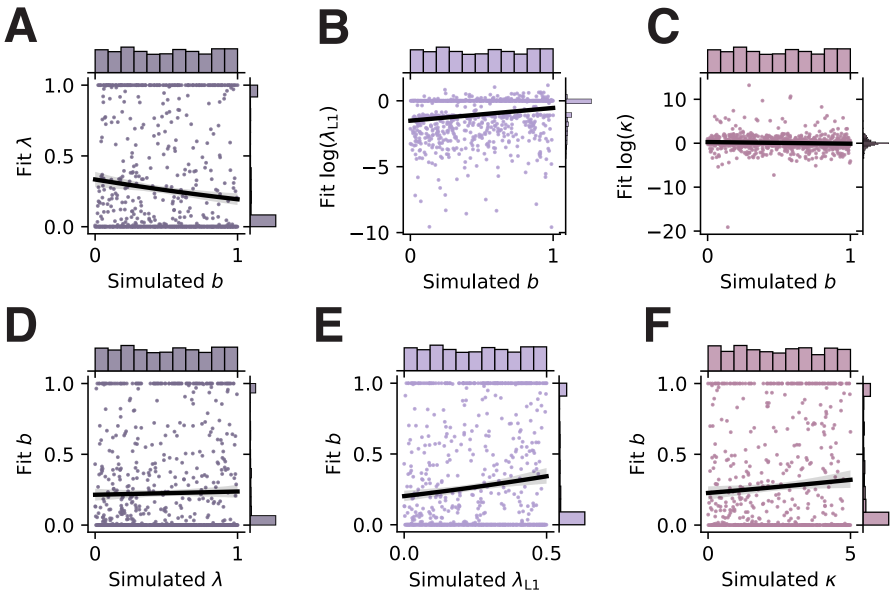


**Figure S9: $b$ recovery within feature-based successor features models.** Relationships between simulated $b$ and fit (**A**) $\lambda$, (**B**) $\lambda_{L1}$, and (**C**) $\kappa$. Relationships between fit $b$ and simulated (**D**) $\lambda$, (**E**) $\lambda_{L1}$, and (**F**) $\kappa$.

### Additional Supplementary Figures

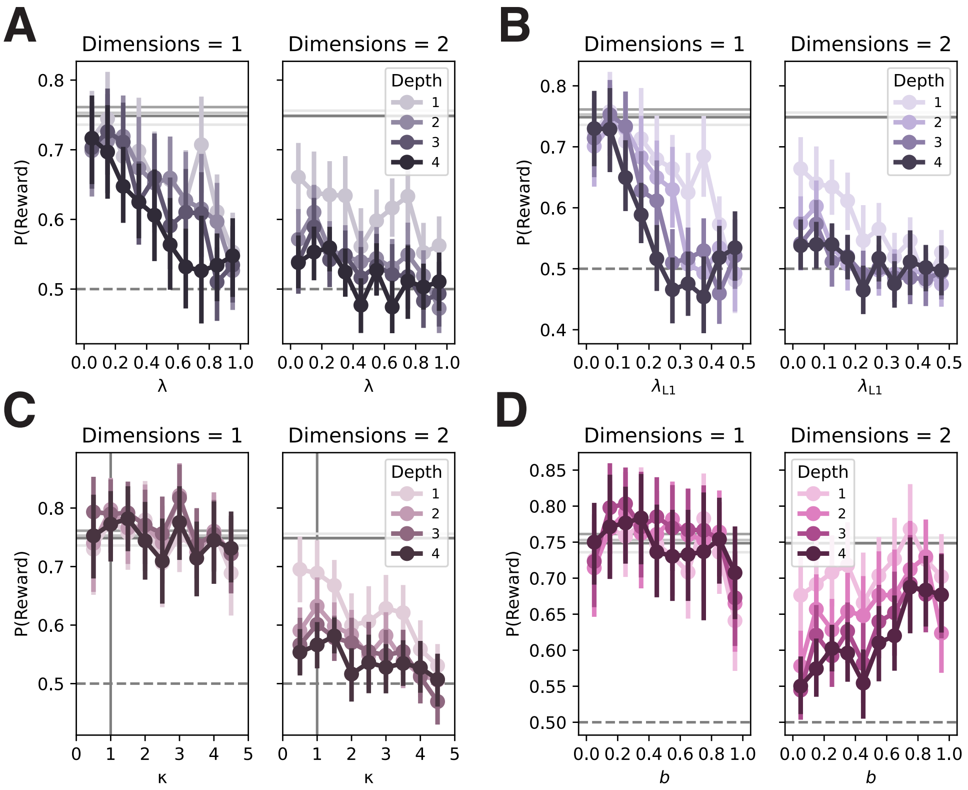


**Figure S10: Feature-based successor features simulations – performance on old test trials.** Average proportion of total reward earned in one- and two-dimensional environments by causal depth by (**A**) decay parameter $\lambda$ in the forgetting model, (**B**) L1 regularization parameter $\lambda_{L1}$ in the L1 regularization model, (**C**) power parameter $\kappa$ in the non-linear value inference model (the vertical line indicates linear inference), and (**D**) causal-bias magnitude parameter in the bias model. Grey lines indicate averages for the causal learner model. None of the unselective bias mechanisms (forgetting, L1 regularization, non-linear value inference) are able to recover the behavioral performance of the selective bias mechanism. See Methods, Computational Models, Feature-based Model for model specifications.

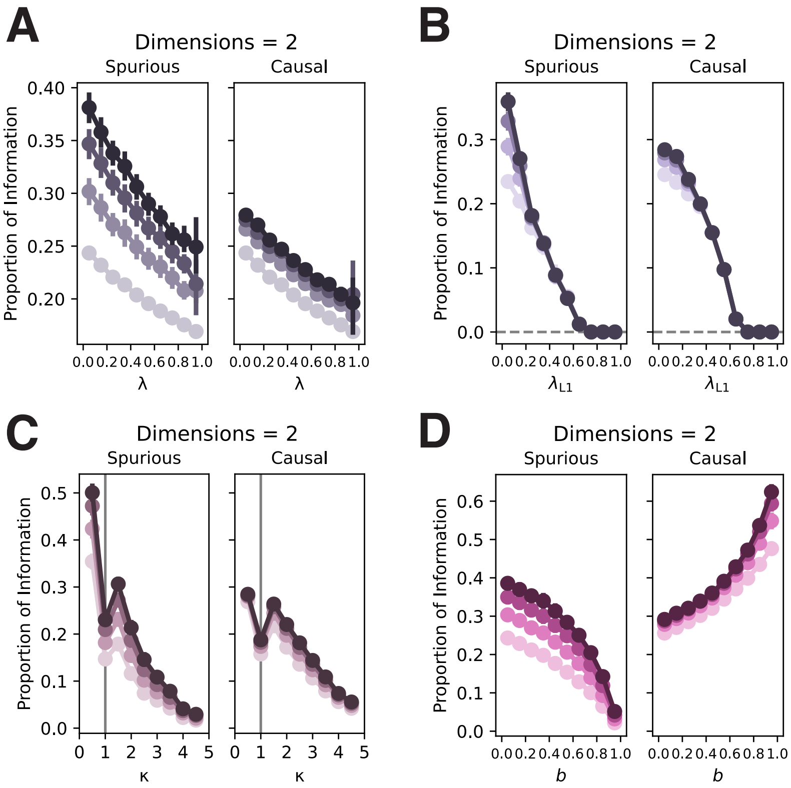


**Figure S11: Feature-based successor features simulations – spurious and causal information at inference.** To identify how spurious and causal information influences inference, we computed target predictions for all single-feature targets (i.e., computing Equation 6, defining $\boldsymbol{w}$ as the identity matrix; note, this definition is different for the non-linear value inference model). We then identify which of these predictions reflect spurious and causal associations, and compute the proportion of spurious/causal information as the sum of the spurious/causal information divided by the total information (spurious + causal + identity). Here, we plot spurious and causal information in the two-dimensional environment by causal depth (darker lines = deeper) and (**A**) decay parameter $\lambda$ in the forgetting model, (**B**) L1 regularization parameter $\lambda_{L1}$ in the L1 regularization model, (**C**) power parameter $\kappa$ in the non-linear value inference model, and (**D**) causal-bias magnitude parameter in the bias model. Only the selective bias mechanism is able to erode spurious information while preserving and even bolstering causal information. See Methods, Computational Models, Feature-based Model for model specifications.

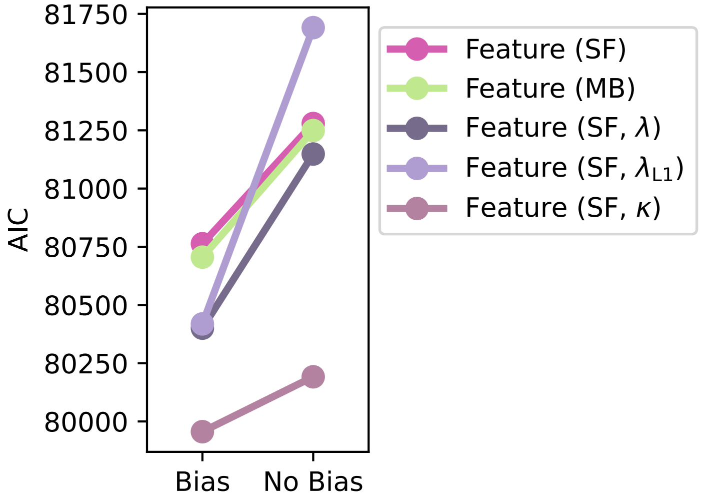


**Figure S12: Group-level feature-based computational model fits, with versus without bias.** Group-level model fits are computed as the sum of AICs by model across participants. Including the bias parameter $b$ improved model fits for all models, despite the AIC metric penalizing for the extra parameter. This indicates both that participants employed a bias on multidimensional predictive learning, and that this selective bias affected learning above and beyond unselective noise reduction mechanisms such as forgetting ($\lambda$), L1 regularization ($\lambda_{L1}$), and non-linear value inference ($\kappa$).

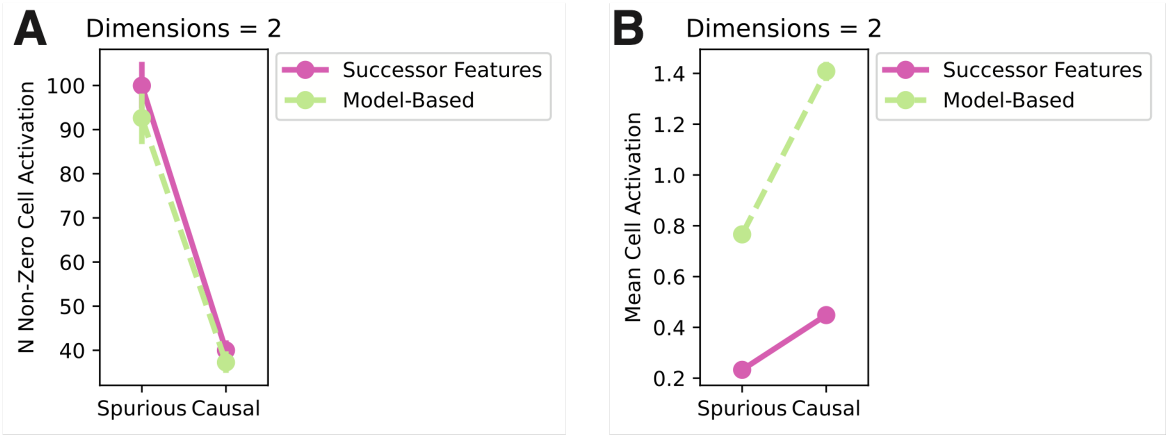


**Figure S13: Causal and spurious information in simulated inferences.** Using simulated agents' learned predictive representations $\boldsymbol{M}$ in the two-dimensional environment (Fig. 2), inferences were computed for each feature in the task independently by calculating Equations 6 and 20 – for the successor features and model-based learner, respectively – defining $\boldsymbol{w}$ as the identity matrix. Note, for the successor features learner, this simply recovers $\boldsymbol{M}$. In the resulting value matrix, we compute the number of cells with non-zero values (**A**) and the mean value across non-zero cells (**B**) for spurious and causal inference, separately. These analyses show that, firstly, spurious information was more diffuse, being spread out over more cells (**A**) while having weaker activations in those cells (**B**). Secondly, model-based inference produced greater mean activation overall for both causal and spurious information (**B**). The likely downstream effect of this greater activation is more deterministic behavior, which is perhaps why we see marginally better model and $b$ recovery for the model-based learner (Fig. S4&7).

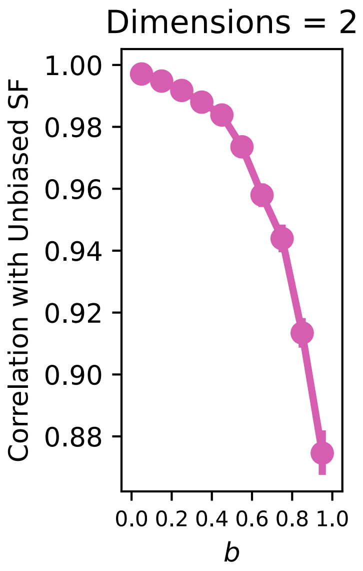


**Figure S14: Correlations between unbiased and biased successor matrices.** Simulated agents' converged successor matrices after learning in two-dimensional environments are compared across degrees of bias $b$ with a standard, unbiased SF learner. As $b$ increases, the successor matrix increasingly diverges from the unbiased successor matrix.

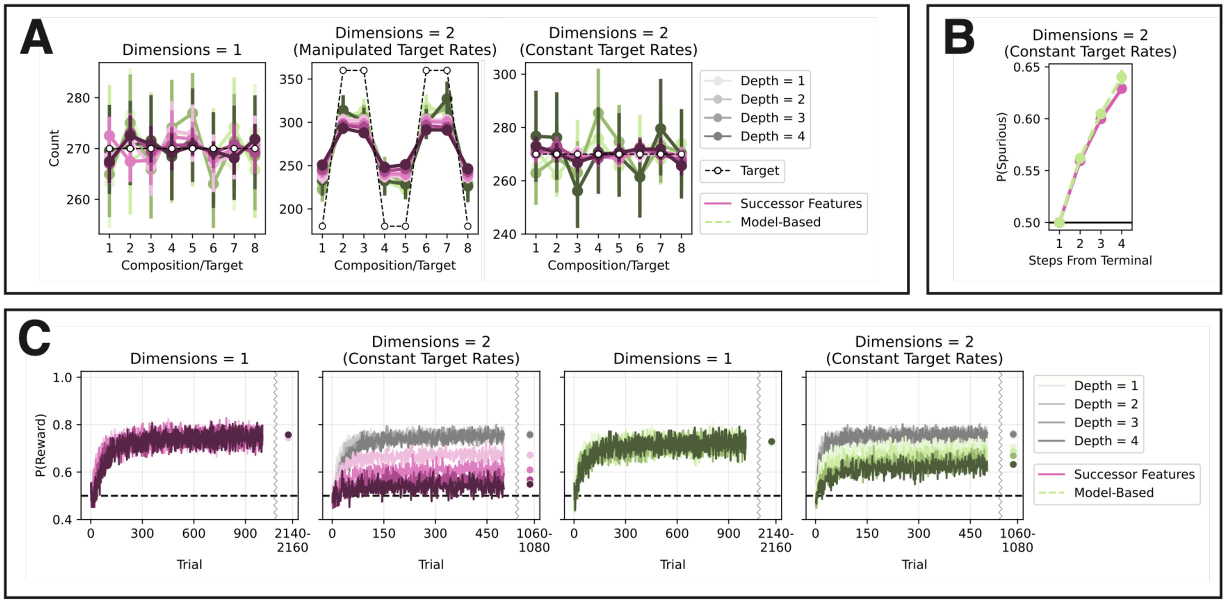


**Figure S15: Simulated learning with constant target frequencies.** (**A**) Average training counts for each possible composition (colored lines) and target (dashed lines) when the feature-based models were simulated in a two-dimensional environment with manipulated target frequencies (Fig. 2, S10&11, S13&14, S16) or constant target frequencies (this Fig. B&C; see Table S2 for specifications). Target frequencies were never manipulated in one-dimensional environments. Composition-target pairs along the x-axis are causally linked. When target frequencies were manipulated, agents adjusted their composition rates accordingly, which in turn would have affected the rates of observed start-successor transitions – this is the basis of the frequency sensitivity analysis (Fig. 2E&3C). (**B&C**) Spurious information distorts predictive learning, even when targets are shown at the same constant rate. (**B**) Proportion of spurious information at inference by model class and the number of steps a feature is from the terminal feature. (**C**) Mean learning curves for the successor features learners (pink lines) and model-based learners (green lines) in one- and two-dimensional environments by causal depth. In the two-dimensional environments, grey lines are causal learners. P(Reward) is the proportion of the maximum reward agents could earn on each trial.

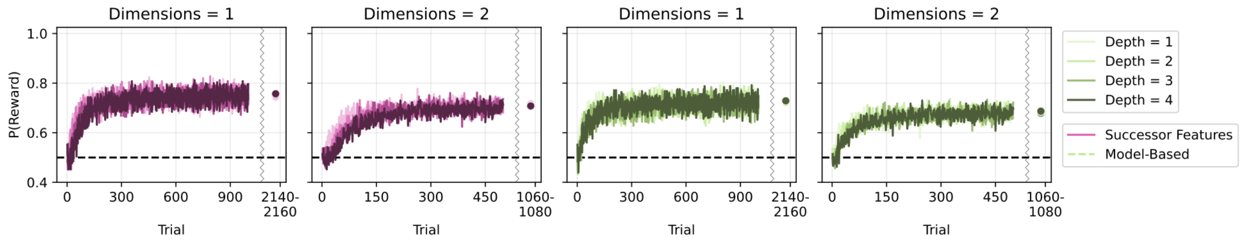


**Figure S16: Training curves for simulated conjunctive learners.** Mean learning curves for the successor features conjunctive learners (pink lines) and model-based conjunctive learners (green lines) in one- and two-dimensional environments by causal depth. In the two-dimensional environments, grey lines are causal learners. P(Reward) is the proportion of the maximum reward agents could earn on each trial. The task and environments were identical to that of the simulations reported in Fig. 2. In one-dimensional environments, feature-based and conjunctive learning are identical processes. In two-dimensional environments, spurious transitions do not arise at the conjunctive level, preventing conjunctive learning from developing the distortions observed for the feature-based learners (Fig. 2).

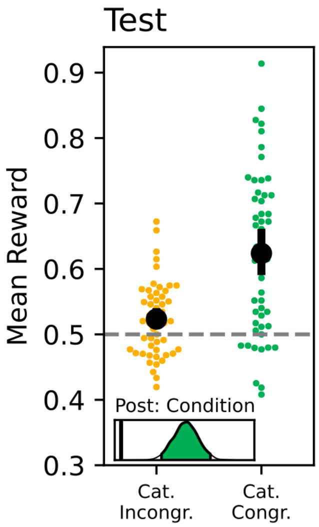


**Figure S17: Test reward earnings by category-congruency condition.** Error bars are 95% HDIs. The density plot shows the posterior of the coefficient for the effect of category-congruency condition, with the black line indicating 0.

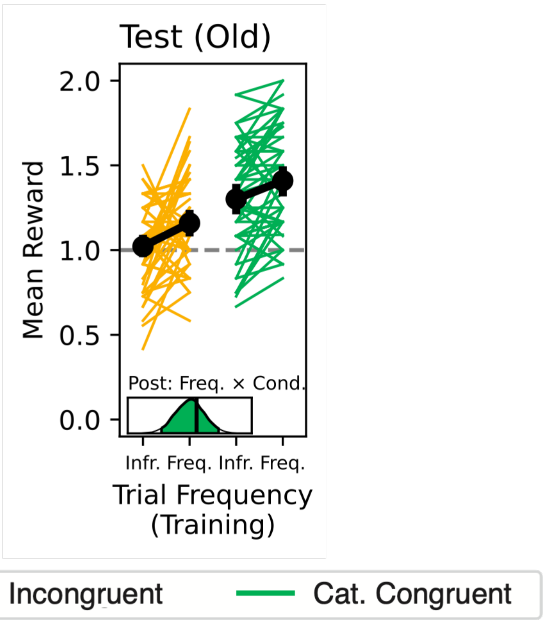


**Figure S18: Test reward earnings on old trials by training target frequency and category-congruency condition.** Error bars are 95% HDIs. The density plot shows the posterior of the coefficient for the interaction between training trial frequency and category-congruency condition, with the black line indicating 0.

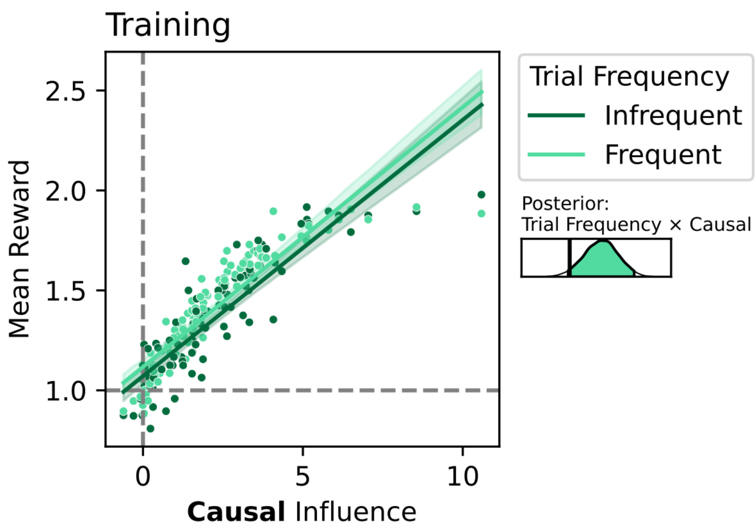


**Figure S19: Training reward earnings by causal transition influence coefficient fit.** Error bars are 95% HDIs. The density plot shows the posterior of the coefficient for the interaction between training trial frequency and causal transition influence, with the black line indicating 0.

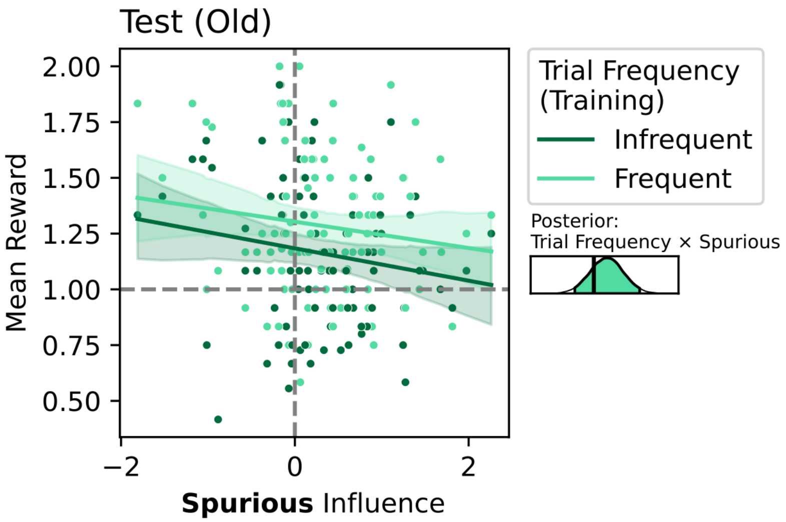


**Figure S20: Test (old) reward earnings by spurious transition influence coefficient fit.** Error bars are 95% HDIs. The density plot shows the posterior of the coefficient for the interaction between training trial frequency and spurious transition influence, with the black line indicating 0.

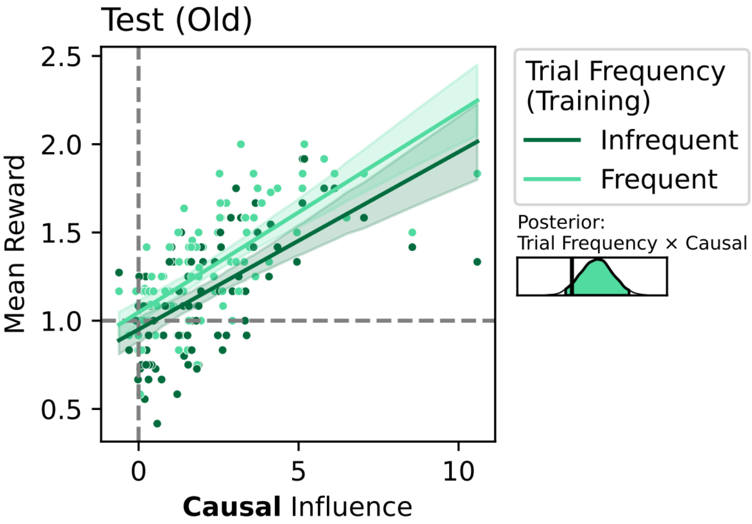


**Figure S21: Test (old) reward earnings by causal transition influence coefficient fit.** Error bars are 95% HDIs. The density plot shows the posterior of the coefficient for the interaction between training trial frequency and causal transition influence, with the black line indicating 0.

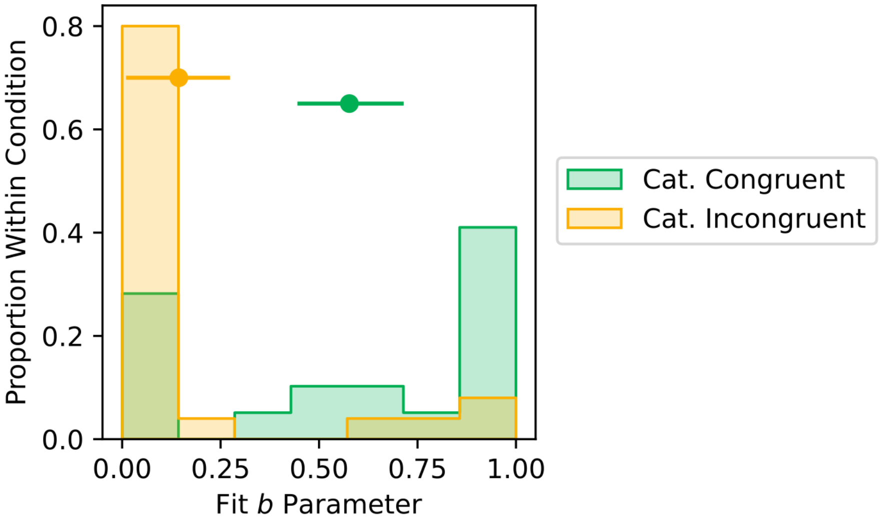


**Figure S22: *b* fit by category-congruency condition.** Error bars are 95% HDIs.

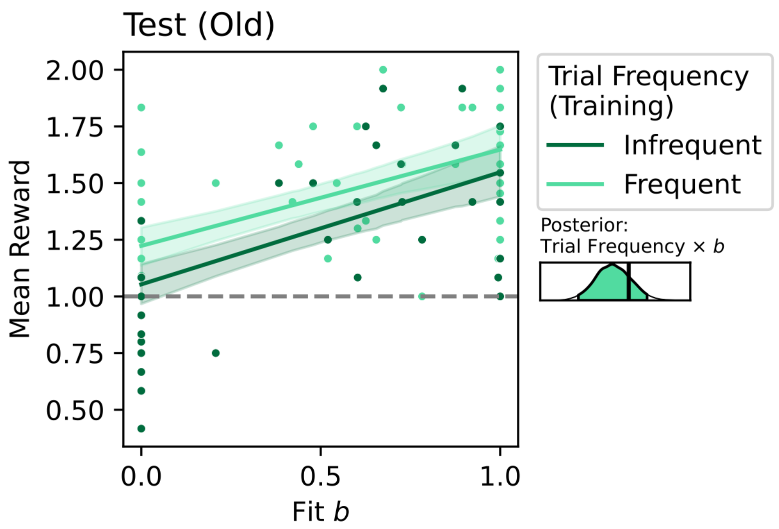


**Figure S23: Test reward earnings on old trials with frequent versus infrequent targets during training by *b* fit.** Error bars are 95% HDIs. The density plot shows the posterior of the coefficient for the interaction between training trial frequency and $b$ fit, with the black line indicating 0.

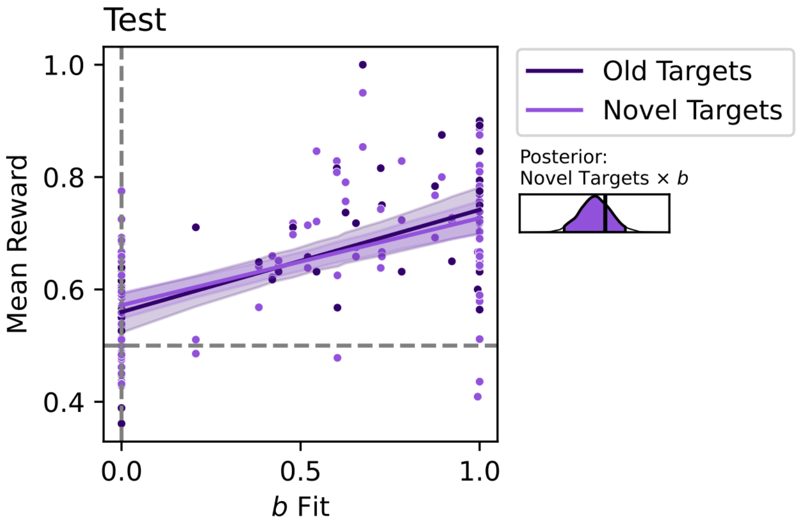


**Figure S24: Test reward earnings on trials with old versus novel targets by $b$ fit.** Error bars are 95% HDIs. The density plot shows the posterior of the coefficient for the interaction between target novelty and $b$ fit, with the black line indicating 0.

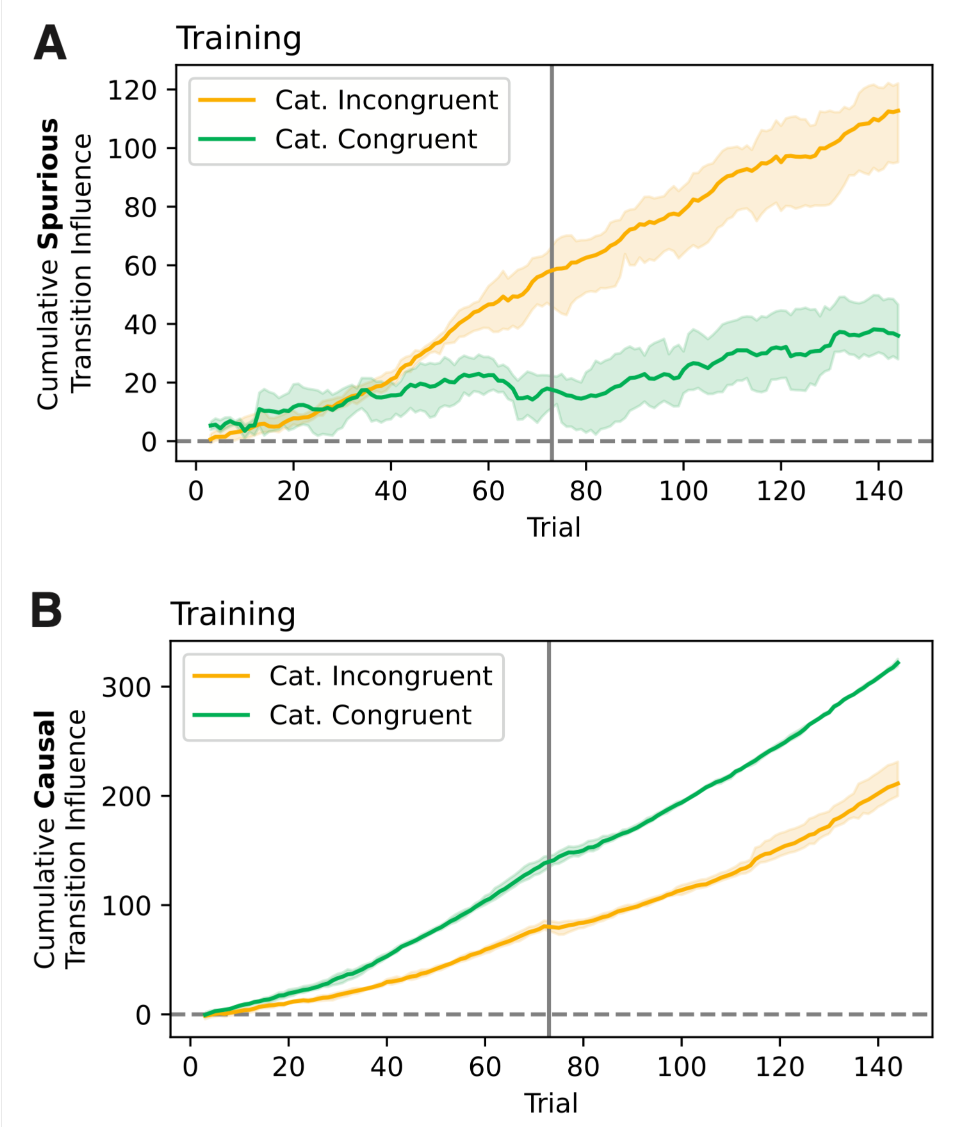


**Figure S25: Cumulative spurious (A) and causal (B) transition influence across training by category-congruency condition.** To explore the trajectory of spurious learning by condition, we ran a trial- rather than agent-wise version of the transition influence analysis. Whereas the preregistered transition influence analysis involved fitting the multinomial logistic regression model separately to each agent's data across training trials (Fig. 3B), here we fit the model separately to each trial's data across agents. This trial-wise analysis was run separately for agents in the category-incongruent and congruent conditions. We then compute the cumulative spurious and causal transition influence coefficients over training trials by condition.

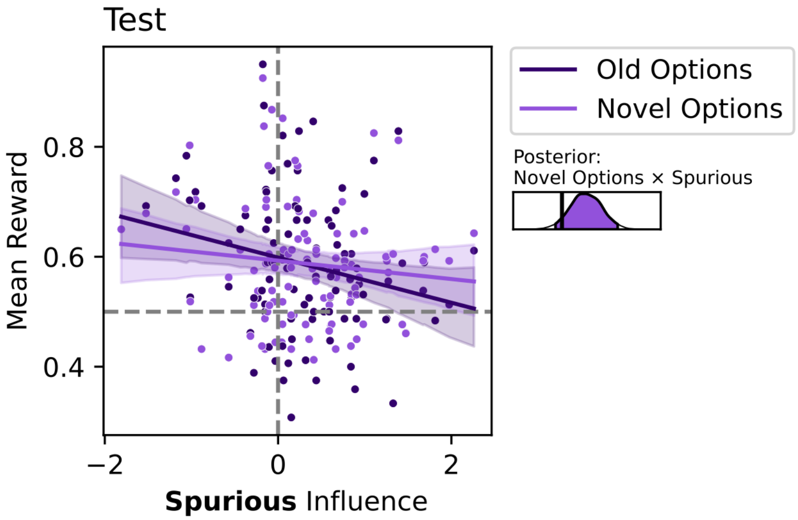


**Figure S26: Test reward earnings on trials with old versus novel options by spurious transition influence coefficient fit.** Error bars are 95% HDIs. The density plot shows the posterior of the coefficient for the interaction between options novelty and spurious transition influence, with the black line indicating 0.

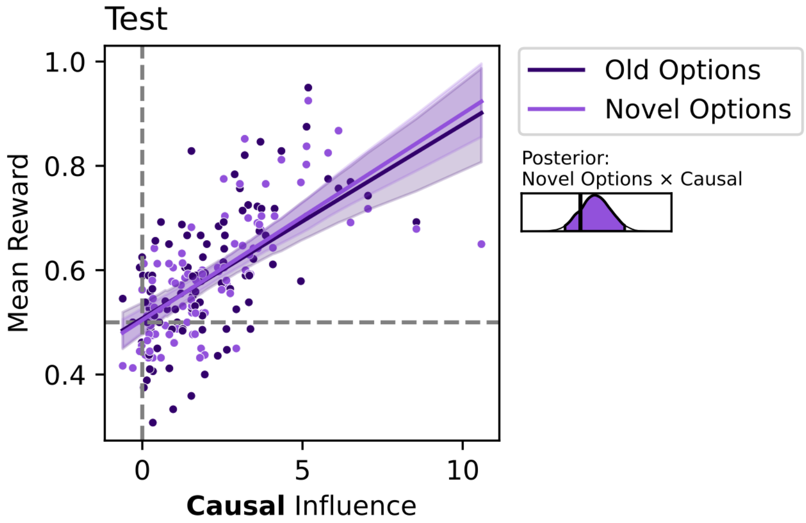


**Figure S27: Test reward earnings on trials with old versus novel options by causal transition influence coefficient fit.** Error bars are 95% HDIs. The density plot shows the posterior of the coefficient for the interaction between options novelty and causal transition influence, with the black line indicating 0.

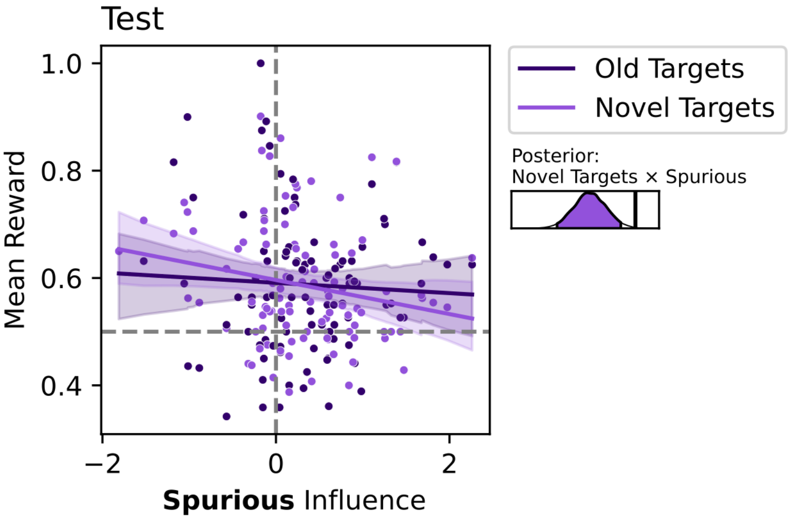


**Figure S28: Test reward earnings on trials with old versus novel targets by spurious transition influence coefficient fit.** Error bars are 95% HDIs. The density plot shows the posterior of the coefficient for the interaction between target novelty and spurious transition influence, with the black line indicating 0.

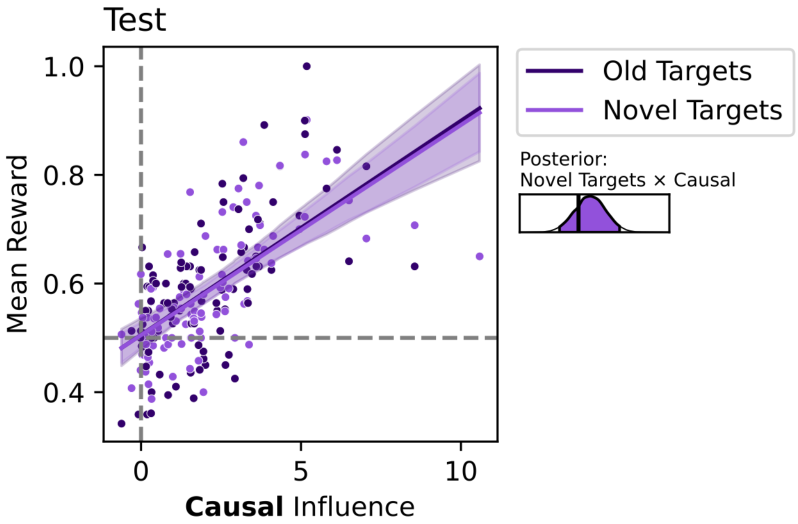


**Figure S29: Test reward earnings on trials with old versus novel targets by causal transition influence coefficient fit.** Error bars are 95% HDIs. The density plot shows the posterior of the coefficient for the interaction between target novelty and causal transition influence, with the black line indicating 0.

### Supplementary Tables

| Causal Depth | | | |
|---|---|---|---|
| 1 | 2 | 3 | 4 |
| A1→A2 | A1→A2→A3 | A1→A2→A3→A4 | A1→A2→A3→A4→A5 |
| A3→A4 | A4→A5→A6 | A5→A6→A7→A8 | A6→A7→A8→A9→A10 |
| B1→B2 | B1→B2→B3 | B1→B2→B3→B4 | B1→B2→B3→B4→B5 |
| B3→B4 | B4→B5→B6 | B5→B6→B7→B8 | B6→B7→B8→B9→B10 |
| C1→C2 | C1→C2→C3 | C1→C2→C3→C4 | C1→C2→C3→C4→C5 |
| C3→C4 | C4→C5→C6 | C5→C6→C7→C8 | C6→C7→C8→C9→C10 |
| D1→D2 | D1→D2→D3 | D1→D2→D3→D4 | D1→D2→D3→D4→D5 |
| D3→D4 | D4→D5→D6 | D5→D6→D7→D8 | D6→D7→D8→D9→D10 |

**Table S1. Causal association definitions – simulations.**

All causal association definitions for 1-, 2-, 3-, and 4-step simulations. All associations are deterministic. Letters correspond to the feature category; numbers correspond to the instance within the category.

| | | Causal Depth | | | |
|---|---|---|---|---|---|
| Manipulated Occurrence Rate | Constant Occurrence Rate (Fig. S15B&C) | 1 | 2 | 3 | 4 |
| Infrequent (x1) | Constant (x1) | {A2, B2} | {A3, B3} | {A4, B4} | {A5, B5} |
| | | {A4, B4} | {A6, B6} | {A8, B8} | {A10, B10} |
| | | {C2, D2} | {C3, D3} | {C4, D4} | {C5, D5} |
| | | {C4, D4} | {C6, D6} | {C8, D8} | {C10, D10} |
| Frequent (x2) | | {A2, B4} | {A3, B6} | {A4, B8} | {A5, B10} |
| | | {A4, B2} | {A6, B3} | {A8, B4} | {A10, B5} |
| | | {C2, D4} | {C3, D6} | {C4, D8} | {C5, D10} |
| | | {C4, D2} | {C6, D3} | {C8, D4} | {C10, D5} |

**Table S2. Target co-occurrence rates – simulations (dimensions = 2).**

Target co-occurrence rates in the two-dimensional environments in the simulations. For most simulations (Fig. 2, Fig. S10&11, S13&14, Fig. S16), half the targets occur twice as frequently as the other half of targets (manipulated occurrence rate). For the simulations in Fig. S15B&C, the target occurrence rate is constant. Letters correspond to the feature category; numbers correspond to the instance within the category.

| | | Online One-Step Experiment | Multi-Step Experiment |
|---|---|---|---|
| Cat. Congruent | Block 1 | A1→A2 | A1→A2→A3 |
| Cat. Congruent | Block 1 | A3→A4 | A4→A5→A6 |
| Cat. Congruent | Block 1 | B1→B2 | B1→B2→B3 |
| Cat. Congruent | Block 1 | B3→B4 | B4→B5→B6 |
| Cat. Congruent | Block 2 | C1→C2 | C1→C2→C3 |
| Cat. Congruent | Block 2 | C3→C4 | C4→C5→C6 |
| Cat. Congruent | Block 2 | D1→D2 | D1→D2→D3 |
| Cat. Congruent | Block 2 | D3→D4 | D4→D5→D6 |
| Cat. Incongruent | Block 1 | C1→A2 | C1→A2→D3 |
| Cat. Incongruent | Block 1 | C3→A4 | C4→A5→D6 |
| Cat. Incongruent | Block 1 | D1→B2 | D1→B2→C3 |
| Cat. Incongruent | Block 1 | D3→B4 | D4→B5→C6 |
| Cat. Incongruent | Block 2 | A1→C2 | A1→C2→B3 |
| Cat. Incongruent | Block 2 | A3→C4 | A4→C5→B6 |
| Cat. Incongruent | Block 2 | B1→D2 | B1→D2→A3 |
| Cat. Incongruent | Block 2 | B3→D4 | B4→D5→A6 |

**Table S3. Causal association definitions – human subjects experiments.**

All causal association definitions in the online pilot one-step experiment and the main multi-step experiment by category-congruency condition and block. Letters correspond to the feature category; numbers correspond to the instance within the category.

| | Occurrence Rate | Online One-Step Experiment | Multi-Step Experiment |
|---|---|---|---|
| Block 1 | Infrequent (x1) | {A2, B2} | {A2, B2}, {A3, B3} |
| | | {A4, B4} | {A5, B5}, {A6, B6} |
| | Frequent (x2) | {A2, B4} | {A2, B5}, {A3, B6} |
| | | {A4, B2} | {A5, B2}, {A6, B3} |
| Block 2 | Infrequent (x1) | {C2, D2} | {C2, D2}, {C3, D3} |
| | | {C4, D4} | {C5, D5}, {C6, D6} |
| | Frequent (x2) | {C2, D4} | {C2, D5}, {C3, D6} |
| | | {C4, D2} | {C5, D2}, {C6, D3} |

**Table S4. Target co-occurrence rates – human subjects experiments.**

Target co-occurrence rates at {step 1} in the pilot online one-step experiment and at {step 1}, {step 2} in the main multi-step experiment. In each block, half the targets occur twice as frequently as the other half of targets. Letters correspond to the feature category; numbers correspond to the instance within the category.

| Variable | M | SD | 95% HDI |
|---|---|---|---|
| Intercept | 0.574 | 0.340 | [-0.104, 1.341] |
| Condition | -0.154 | 0.247 | [-0.626, 0.338] |
| Transition Type | 1.139 | 0.250 | [0.649, 1.615] |
| Condition × Transition Type | 0.875 | 0.352 | [0.173, 1.563] |
| $\sigma$ | 2.195 | 0.065 | [2.073, 2.321] |

**Table S5. Fixed effects for transition influence analysis (initial one-step experiment).**

Linear regression formula:
(transition influence) ~ (condition) + (transition type) + (condition × transition type) + (1|logit comparison)

Linear regression variables:

- transition influence: fit transition influence coefficient
- condition: category-congruency condition (0 = category-incongruent, 1 = category-congruent)
- transition type: transition influence type (0 = spurious, 1 = causal)
- logit comparison: whether the coefficient was from the logit comparing $\frac{P(C=1)}{P(C=0)}$, $\frac{P(C=2)}{P(C=0)}$, or $\frac{P(C=3)}{P(C=0)}$ (random intercepts)

MCMC settings:

- Chains: 4
- Total iterations per chain: 6000
- Warm-up iterations per chain: 4000
- Posterior samples per chain: 2000

| Variable | M | SD | 95% HDI |
|---|---|---|---|
| Intercept | 0.584 | 0.196 | [0.185, 0.963] |
| Condition | -0.555 | 0.215 | [-0.980, -0.136] |
| Transition Type | 0.862 | 0.136 | [0.590, 1.124] |
| Condition × Transition Type | 1.847 | 0.193 | [1.464, 2.225] |
| $\sigma$ | 1.205 | 0.037 | [1.133, 1.278] |

**Table S6. Fixed effects for transition influence analysis.**

Linear regression formula:
(transition influence) ~ (condition) + (transition type) + (condition × transition type) + (1|logit comparison) + (1|subject)

Linear regression variables:

- transition influence: fit transition influence coefficient
- condition: category-congruency condition (0 = category-incongruent, 1 = category-congruent)
- transition type: transition influence type (0 = spurious, 1 = causal)
- logit comparison: whether the coefficient was from the logit comparing $\frac{P(C=1)}{P(C=0)}$, $\frac{P(C=2)}{P(C=0)}$, or $\frac{P(C=3)}{P(C=0)}$ (random intercepts)
- subject: participant identity (random intercepts)

MCMC settings:

- Chains: 4
- Total iterations per chain: 6000
- Warm-up iterations per chain: 4000
- Posterior samples per chain: 2000

| Variable | M | SD | 95% HDI |
|---|---|---|---|
| Intercept | 1.302 | 0.037 | [1.228, 1.372] |
| Condition | 0.143 | 0.053 | [0.040, 0.249] |
| $\sigma$ | 0.677 | 0.004 | [0.669, 0.684] |

**Table S7. Fixed effects for training reward earnings by condition analysis.**

Linear regression formula:
(reward) ~ (condition) + (1|subject)

Linear regression variables:

- reward: reward on training trials
- condition: category-congruency condition (0 = category-incongruent, 1 = category-congruent)
- subject: participant identity (random intercepts)

MCMC settings:

- Chains: 4
- Total iterations per chain: 4000
- Warm-up iterations per chain: 2000
- Posterior samples per chain: 2000

| Variable | M | SD | 95% HDI |
|---|---|---|---|
| Intercept | 0.523 | 0.014 | [0.497, 0.550] |
| Condition | 0.100 | 0.019 | [0.063, 0.139] |
| $\sigma$ | 0.589 | 0.003 | [0.582, 0.595] |

**Table S8. Fixed effects for test reward earnings by condition analysis.**

Linear regression formula:
(reward) ~ (condition) + (1|subject)

Linear regression variables:

- reward: reward on test trials
- condition: category-congruency condition (0 = category-incongruent, 1 = category-congruent)
- subject: participant identity (random intercepts)

MCMC settings:

- Chains: 4
- Total iterations per chain: 4000
- Warm-up iterations per chain: 2000
- Posterior samples per chain: 2000

| Variable | M | SD | 95% HDI |
|---|---|---|---|
| Intercept | 0.501 | 0.012 | [0.476, 0.525] |
| Condition | 0.052 | 0.018 | [0.018, 0.088] |
| $\sigma$ | 0.495 | 0.005 | [0.486, 0.504] |

**Table S9. Fixed effects for reward earnings on two-step test trials by condition analysis.**

Linear regression formula:
(reward) ~ (condition) + (1|subject)

Linear regression variables:

- reward: reward on two-step test trials
- condition: category-congruency condition (0 = category-incongruent, 1 = category-congruent)
- subject: participant identity (random intercepts)

MCMC settings:

- Chains: 4
- Total iterations per chain: 4000
- Warm-up iterations per chain: 2000
- Posterior samples per chain: 2000

| Variable | M | SD | 95% HDI |
|---|---|---|---|
| Intercept | 0.497 | 0.032 | [0.436, 0.562] |
| Condition | 0.275 | 0.045 | [0.188, 0.368] |
| Step | 0.055 | 0.017 | [0.021, 0.087] |
| Condition × Step | -0.094 | 0.023 | [-0.139, -0.047] |
| $\sigma$ | 0.454 | 0.004 | [0.446, 0.462] |

**Table S10. Fixed effects for one-step test reward earnings on 1st vs 2nd step trials by condition.**

Linear regression formula:
(reward) ~ (condition) + (step) + (condition × step) + (1|subject)

Linear regression variables:

- reward: one-step test trial reward
- condition: category-congruency condition (0 = category-incongruent, 1 = category-congruent)
- step: Options step in the causal chains (1 = 1st step, 2 = 2nd step)
- subject: participant identity (random intercepts)

MCMC settings:

- Chains: 4
- Total iterations per chain: 4000
- Warm-up iterations per chain: 2000
- Posterior samples per chain: 2000

| Variable | M | SD | 95% HDI |
|---|---|---|---|
| Intercept | 1.249 | 0.039 | [1.171, 1.324] |
| Condition | 0.182 | 0.056 | [0.072, 0.291] |
| Frequency | 0.081 | 0.017 | [0.048, 0.114] |
| Condition × Frequency | -0.063 | 0.024 | [-0.110, -0.016] |
| $\sigma$ | 0.676 | 0.004 | [0.668, 0.684] |

**Table S11. Fixed effects for training frequency sensitivity by condition analysis.**

Linear regression formula:
(reward) ~ (condition) + (frequency) + (condition × frequency) + (1|subject)

Linear regression variables:

- reward: training trial reward
- condition: category-congruency condition (0 = category-incongruent, 1 = category-congruent)
- frequency: target frequency during training (0 = infrequent, 1 = frequent)
- subject: participant identity (random intercepts)

MCMC settings:

- Chains: 4
- Total iterations per chain: 4000
- Warm-up iterations per chain: 2000
- Posterior samples per chain: 2000

| Variable | M | SD | 95% HDI |
| --- | --- | --- | --- |
| Intercept | 1.024 | 0.041 | [0.943, 1.103] |
| Condition | 0.279 | 0.057 | [0.171, 0.392] |
| Frequency | 0.134 | 0.039 | [0.057, 0.211] |
| Condition × Frequency | -0.027 | 0.056 | [-0.135, 0.085] |
| $\sigma$ | 0.688 | 0.010 | [0.668, 0.707] |

**Table S12. Fixed effects for test frequency sensitivity by condition analysis.**

Linear regression formula:
(reward) ~ (condition) + (frequency) + (condition × frequency) + (1|subject)

Linear regression variables:

- reward: test trial reward
- condition: category-congruency condition (0 = category-incongruent, 1 = category-congruent)
- frequency: target frequency during training (0 = infrequent, 1 = frequent)
- subject: participant identity (random intercepts)

MCMC settings:

- Chains: 4
- Total iterations per chain: 4000
- Warm-up iterations per chain: 2000
- Posterior samples per chain: 2000

| Variable | M | SD | 95% HDI |
|---|---|---|---|
| Intercept | 1.212 | 0.028 | [1.159, 1.268] |
| Causal Influence | 0.067 | 0.010 | [0.048, 0.087] |
| Spurious Influence | -0.043 | 0.014 | [-0.071, -0.017] |
| Frequency | 0.020 | 0.011 | [-0.001, 0.042] |
| Causal Influence × Spurious Influence | 0.003 | 0.003 | [-0.003, 0.008] |
| Causal Influence × Frequency | 0.007 | 0.004 | [-0.0003, 0.014] |
| Spurious Influence × Frequency | 0.047 | 0.013 | [0.023, 0.073] |
| Causal Influence × Spurious Influence × Frequency | 0.002 | 0.003 | [-0.003, 0.008] |
| $\sigma$ | 0.675 | 0.002 | [0.670, 0.679] |

**Table S13. Fixed effects for training frequency sensitivity by transition influence analysis.**

Linear regression formula:
(reward) ~ (causal influence) + (spurious influence) + (frequency) + (causal influence × spurious influence) + (causal influence × frequency) + (spurious influence × frequency) + (causal influence × spurious influence × frequency) + (1|subject) + (1|logit comparison)

Linear regression variables:

- reward: training trial reward
- causal influence: fit causal transition influence coefficient
- spurious influence: fit spurious transition influence coefficient
- frequency: target frequency during training (0 = infrequent, 1 = frequent)
- subject: participant identity (random intercepts)
- logit comparison: whether the coefficient was from the logit comparing $\frac{P(C=1)}{P(C=0)}$, $\frac{P(C=2)}{P(C=0)}$, or $\frac{P(C=3)}{P(C=0)}$ (random intercepts)

MCMC settings:

- Chains: 4
- Total iterations per chain: 4000
- Warm-up iterations per chain: 2000
- Posterior samples per chain: 2000

| Variable | M | SD | 95% HDI |
|---|---|---|---|
| Intercept | 0.995 | 0.036 | [0.923, 1.063] |
| Causal Influence | 0.086 | 0.010 | [0.065, 0.106] |
| Spurious Influence | -0.054 | 0.028 | [-0.108, 0.001] |
| Frequency | 0.088 | 0.025 | [0.038, 0.136] |
| Causal Influence × Spurious Influence | 0.012 | 0.006 | [0.000, 0.022] |
| Causal Influence × Frequency | 0.013 | 0.008 | [-0.003, 0.030] |
| Spurious Influence × Frequency | 0.023 | 0.030 | [-0.035, 0.083] |
| Causal Influence × Spurious Influence × Frequency | -0.002 | 0.006 | [-0.014, 0.010] |
| $\sigma$ | 0.679 | 0.006 | [0.667, 0.689] |

**Table S14. Fixed effects for test frequency sensitivity by transition influence analysis.**

Linear regression formula:
(reward) ~ (causal influence) + (spurious influence) + (frequency) + (causal influence × spurious influence) + (causal influence × frequency) + (spurious influence × frequency) + (causal influence × spurious influence × frequency) + (1|subject) + (1|logit comparison)

Linear regression variables:

- reward: test trial reward
- causal influence: fit causal transition influence coefficient
- spurious influence: fit spurious transition influence coefficient
- frequency: target frequency during training (0 = infrequent, 1 = frequent)
- subject: participant identity (random intercepts)
- logit comparison: whether the coefficient was from the logit comparing $\frac{P(C=1)}{P(C=0)}$, $\frac{P(C=2)}{P(C=0)}$, or $\frac{P(C=3)}{P(C=0)}$ (random intercepts)

MCMC settings:

- Chains: 4
- Total iterations per chain: 8000
- Warm-up iterations per chain: 4000
- Posterior samples per chain: 4000

| Variable | M | SD | 95% HDI |
|---|---|---|---|
| Intercept | 0.550 | 0.022 | [0.509, 0.595] |
| Novel Options | -0.005 | 0.015 | [-0.034, 0.024] |
| Novel Target | -0.010 | 0.015 | [-0.038, 0.020] |
| Condition | 0.071 | 0.032 | [0.011, 0.135] |
| Novel Options × Condition | 0.014 | 0.021 | [-0.028, 0.056] |
| Novel Target × Condition | 0.018 | 0.021 | [-0.024, 0.059] |
| $\sigma$ | 0.483 | 0.003 | [0.476, 0.489] |

**Table S15. Fixed effects for generalization by condition analysis.**

Linear regression formula:
(reward) ~ (novel options) + (novel target) + (condition) + (novel options × condition) + (novel target × condition) + (1|subject)

Linear regression variables:

- reward: test trial reward
- novel options: options novelty (old = 0, novel = 1)
- novel target: target novelty (old = 0, novel = 1)
- condition: category-congruency condition (0 = category-incongruent, 1 = category-congruent)
- subject: participant identity (random intercepts)

MCMC settings:

- Chains: 4
- Total iterations per chain: 4000
- Warm-up iterations per chain: 2000
- Posterior samples per chain: 2000

| Variable | M | SD | 95% HDI |
|---|---|---|---|
| Intercept | 0.953 | 0.027 | [0.898, 1.005] |
| Condition | -0.018 | 0.038 | [-0.090, 0.059] |
| $\sigma$ | 0.713 | 0.013 | [0.686, 0.738] |

**Table S16. Fixed effects for composition spurious predictiveness by condition analysis.**

Linear regression formula:
(spurious predictiveness) ~ (condition) + (1|subject)

Linear regression variables:

- spurious predictiveness: the spurious predictiveness of the test trial's composition
- condition: category-congruency condition (0 = category-incongruent, 1 = category-congruent)
- subject: participant identity (random intercepts)

MCMC settings:

- Chains: 4
- Total iterations per chain: 4000
- Warm-up iterations per chain: 2000
- Posterior samples per chain: 2000

| Variable | M | SD | 95% HDI |
|---|---|---|---|
| Intercept | 0.560 | 0.192 | [0.125, 0.939] |
| $b$ | -0.752 | 0.214 | [-1.175, -0.336] |
| Transition Type | 0.732 | 0.183 | [0.372, 1.095] |
| $b$ × Transition Type | 4.178 | 0.308 | [3.555, 4.755] |
| $\sigma$ | 1.279 | 0.047 | [1.192, 1.373] |

**Table S17. Fixed effects for transition influence by $b$ analysis.**

Linear regression formula:
(transition influence) ~ ($b$) + (transition type) + ($b$ × transition type) + (1|logit comparison)

Linear regression variables:

- transition influence: fit transition influence coefficient
- $b$: fit causal-bias parameter in the feature-based model
- transition type: transition influence type (0 = spurious, 1 = causal)
- logit comparison: whether the coefficient was from the logit comparing $\frac{P(C=1)}{P(C=0)}$, $\frac{P(C=2)}{P(C=0)}$, or $\frac{P(C=3)}{P(C=0)}$ (random intercepts)

MCMC settings:

- Chains: 4
- Total iterations per chain: 4000
- Warm-up iterations per chain: 2000
- Posterior samples per chain: 2000

| Variable | M | SD | 95% HDI |
|---|---|---|---|
| Intercept | 1.222 | 0.033 | [1.157, 1.284] |
| $b$ | 0.494 | 0.056 | [0.382, 0.604] |
| Frequency | 0.071 | 0.020 | [0.033, 0.111] |
| $b$ × Frequency | -0.058 | 0.033 | [-0.126, 0.004] |
| $\sigma$ | 0.647 | 0.005 | [0.638, 0.656] |

**Table S18. Fixed effects for training frequency sensitivity by $b$ analysis.**

Linear regression formula:
(reward) ~ ($b$) + (frequency) + ($b$ × frequency) + (1|subject)

Linear regression variables:

- reward: training trial reward
- $b$: fit causal-bias parameter in the feature-based model
- frequency: target frequency during training (0 = infrequent, 1 = frequent)
- subject: participant identity (random intercepts)

MCMC settings:

- Chains: 4
- Total iterations per chain: 4000
- Warm-up iterations per chain: 2000
- Posterior samples per chain: 2000

| Variable | M | SD | 95% HDI |
|---|---|---|---|
| Intercept | 1.054 | 0.043 | [0.973, 1.143] |
| $b$ | 0.494 | 0.074 | [0.354, 0.644] |
| Frequency | 0.169 | 0.046 | [0.075, 0.257] |
| $b$ × Frequency | -0.069 | 0.078 | [-0.223, 0.081] |
| $\sigma$ | 0.652 | 0.012 | [0.629, 0.675] |

**Table S19. Fixed effects for test frequency sensitivity by $b$ analysis.**

Linear regression formula:
(reward) ~ ($b$) + (frequency) + ($b$ × frequency) + (1|subject)

Linear regression variables:

- reward: test trial reward
- $b$: fit causal-bias parameter in the feature-based model
- frequency: target frequency during training (0 = infrequent, 1 = frequent)
- subject: participant identity (random intercepts)

MCMC settings:

- Chains: 4
- Total iterations per chain: 4000
- Warm-up iterations per chain: 2000
- Posterior samples per chain: 2000

| Variable | M | SD | 95% HDI |
|---|---|---|---|
| Intercept | 0.570 | 0.025 | [0.523, 0.620] |
| Novel Options | -0.011 | 0.018 | [-0.045, 0.024] |
| Novel Target | 0.007 | 0.018 | [-0.030, 0.042] |
| $b$ | 0.169 | 0.042 | [0.093, 0.258] |
| Novel Options × $b$ | 0.011 | 0.031 | [-0.046, 0.073] |
| Novel Target × $b$ | -0.020 | 0.030 | [-0.079, 0.039] |
| $\sigma$ | 0.473 | 0.004 | [0.465, 0.480] |

**Table S20. Fixed effects for generalization by $b$ analysis.**

Linear regression formula:
(reward) ~ (novel options) + (novel target) + ($b$) + (novel options × $b$) + (novel target × $b$) + (1|subject)

Linear regression variables:

- reward: test trial reward
- novel options: options novelty (old = 0, novel = 1)
- novel target: target novelty (old = 0, novel = 1)
- $b$: fit causal-bias parameter in the feature-based model
- subject: participant identity (random intercepts)

MCMC settings:

- Chains: 4
- Total iterations per chain: 4000
- Warm-up iterations per chain: 2000
- Posterior samples per chain: 2000

| Variable | M | SD | 95% HDI |
|---|---|---|---|
| Intercept | 0.940 | 0.034 | [0.870, 1.005] |
| $b$ | -0.020 | 0.057 | [-0.129, 0.091] |
| $\sigma$ | 0.717 | 0.017 | [0.684, 0.751] |

**Table S21. Fixed effects for composition spurious predictiveness by $b$ analysis.**

Linear regression formula:
(spurious predictiveness) ~ ($b$) + (1|subject)

Linear regression variables:

- spurious predictiveness: the spurious predictiveness of the test trial's composition
- $b$: fit causal-bias parameter in the feature-based model
- subject: participant identity (random intercepts)

MCMC settings:

- Chains: 4
- Total iterations per chain: 4000
- Warm-up iterations per chain: 2000
- Posterior samples per chain: 2000

| Model | Group AIC | P(Best Fit) | Relative Likelihoods of Best Fits |
|---|---|---|---|
| **Feature** (MB) | 81249.10 | 0.069 | 0.4729 (0.1321) |
| **Feature** (MB, Bias) | 80705.40 | 0.039 | 0.7768 (0.2203) |
| **Feature** (SF) | 81279.13 | 0.049 | 0.2380 (0.0389) |
| **Feature** (SF, Bias) | 80762.99 | 0 | - |
| **Feature** (SF, Decay) | 81148.22 | 0.020 | 0.6620 (0.4508) |
| **Feature** (SF, Bias, Decay) | 80400.45 | 0.039 | 0.6053 (0.2145) |
| **Feature** (SF, L1 Reg.) | 81690.90 | 0 | - |
| **Feature** (SF, Bias, L1 Reg.) | 80418.74 | 0.147 | 0.8203 (0.2439) |
| **Feature** (SF, Non-Lin.) | 80191.66 | 0.167 | 0.5765 (0.1561) |
| **Feature** (SF, Bias, Non-Lin.) | 79956.01 | 0.098 | 0.5569 (0.1931) |
| **Conj.** (MB) | 81477.43 | 0.167 | 0.8218 (0.2436) |
| **Conj.** (SF) | 82383.84 | 0.059 | 0.7550 (0.2023) |
| **Conj.** (MB, Sampler) | 82702.01 | 0.020 | 0.6229 (0.2981) |
| **Conj.** (SF, Sampler) | 83274.88 | 0.049 | 0.6475 (0.2751) |
| **Null** | 89330.04 | 0.078 | 0.3594 (0.1126) |

**Table S22.** Computational model fits. Relative likelihoods are the means (standard deviations) for models that provided a best fit to a participant's data. A higher value for a model reflects that the model provided a particularly effective account of behavior for participants who were best fit by it.

| Model | $\alpha$ | $\beta$ | $b$ | $\lambda$ | $\lambda_{\mathrm{L1}}$ | $\kappa$ | $\omega^{SIM}$ | $\psi$ |
|---|---|---|---|---|---|---|---|---|
| **Feature** (MB) | 0.153 (0.174) | 0.548 (1.344) | - | - | - | - | - | - |
| **Feature** (MB, Bias) | 0.011 (0.009) | 3.889 (1.469) | 0.819 (0.211) | - | - | - | - | - |
| **Feature** (SF) | 0.164 (0.179) | 1.187 (1.272) | - | - | - | - | - | - |
| **Feature** (SF, Bias) | NA | NA | NA | - | - | - | - | - |
| **Feature** (SF, Decay) | 0.985 (0.021) | -1.168 (0.244) | - | 0.938 (0.088) | - | - | - | - |
| **Feature** (SF, Bias, Decay) | 0.401 (0.246) | 2.009 (0.515) | 0.677 (0.257) | 0.862 (0.162) | - | - | - | - |
| **Feature** (SF, L1 Reg.) | NA | NA | - | - | NA | - | - | - |
| **Feature** (SF, Bias, L1 Reg.) | 0.604 (0.229) | 1.863 (0.358) | 0.828 (0.239) | - | 0.273 (0.054) | - | - | - |
| **Feature** (SF, Non-Lin.) | 0.270 (0.296) | 2.436 (2.521) | - | - | - | -8.134 (10.03) | - | - |
| **Feature** (SF, Bias, Non-Lin.) | 0.481 (0.340) | 2.171 (2.444) | 0.766 (0.261) | - | - | 0.448 (2.789) | - | - |
| **Conj.** (MB) | 0.653 (0.281) | 0.377 (0.376) | - | - | - | - | - | - |
| **Conj.** (SF) | 0.556 (0.431) | 1.362 (2.713) | - | - | - | - | - | - |
| **Conj.** (MB, Sampler) | 0.050 (0.070) | 3.315 (3.563) | - | - | - | - | 0.119 (0.151) | 1.009 (1.427) |
| **Conj.** (SF, Sampler) | 0.086 (0.120) | 4.742 (2.523) | - | - | - | - | 0.424 (0.280) | 0.418 (0.345) |
| **Null** | - | - | - | - | - | - | - | - |

**Table S23**. Computational model parameter fits. Parameter values are the means (standard deviations) across participants best fit by the model. The $\beta$, $\kappa$, and $\psi$ parameters are log transformed. NA indicates no participants were best fit by the model.
